# Ciz1 safeguards *Drosophila* wing development by suppressing oxidative stress

**DOI:** 10.64898/2026.06.21.733590

**Authors:** Xiaojiao Li, Cuiping Wang, Yuanyuan Zhang, Huan Liu, Min Hou, Xiuxiu Liu, Yuanlin Su, Yanqin Gong, Huiyuan Ding, Qiao Liu, Yaoqin Gong, Gongping Sun

**Affiliations:** Key Laboratory for Experimental Teratology of Ministry of Education, Shandong Key Laboratory of Mental Disorders and Intelligent Control, Department of Histology and Embryology, School of Basic Medical Sciences, Cheeloo College of Medicine, Shandong University, Jinan, Shandong, 250012, China; Key Laboratory for Experimental Teratology of Ministry of Education, and Department of Genetics, School of Basic Medical Sciences, Cheeloo College of Medicine, Shandong University, Jinan 250012, China

**Keywords:** Ciz1, *Drosophila* wing, oxidative stress, JNK, EGFR

## Abstract

Cell proliferation and fate specification are fundamental processes that ensure the generation of organs with proper size and patterning. Oxidative stress caused by accumulation of reactive oxygen species (ROS) can lead to cell cycle arrest, senescence, cell death and cell fate misspecification, thereby impairing normal development and contributing to many pathological processes. In this study, we identify *Drosophila* Ciz1 as a critical factor that safeguards epithelial homeostasis and development by preventing oxidative stress. Knockdown of *Ciz1* in the *Drosophila* wing imaginal disc, an epithelial tissue that serves as the larval precursor of the adult wing, results in a small wing phenotype accompanied by thickened and ectopic veins. We further demonstrate that reduced Ciz1 expression leads to accumulation of donut-shaped mitochondria and elevated ROS levels. The increased oxidative stress subsequently suppresses proliferation via activation of JNK and promotes excessive vein formation by upregulating Rhomboid, a positive regulator of EGFR signaling. Interestingly, although Ciz1 is a zinc finger protein that predominantly localizes to the nucleus, neither its zinc finger motifs nor its nuclear localization is required for suppression of oxidative stress. Instead, the prion-like domain in its N-terminal part is essential for this activity. Our work identifies Ciz1 as an important factor in preventing oxidative stress and maintaining epithelial homeostasis.

## Introduction

Reactive oxygen species (ROS) are primarily generated in mitochondria as byproducts of oxidative phosphorylation. At moderate levels, ROS serve as important signaling molecules that can promote cell survival, proliferation and differentiation, thereby facilitating tissue development, wound healing and regeneration(1). However, high levels of ROS can cause oxidative damage to cellular components such as lipids, proteins and nucleic acids, leading to cell cycle arrest, senescence and cell death. Increased oxidative stress has been implicated in aging, cancer, cardiovascular diseases, neurodegenerative disorders, diabetes, arthritis, and other pathogenic conditions(2, 3). To counteract ROS accumulation, cells establish an antioxidant system that includes enzymatic oxidants (superoxide dismutase, catalase, glutathione peroxidase, peroxiredoxin and thioredoxin) and non-enzymatic oxidants (coenzyme Q10, melatonin, α-lipoic acid and uric acid)(1–3).

Cell proliferation and fate specification are fundamental processes that ensure the generation of organs with proper size and patterning. The *Drosophila* wing imaginal disc is a larval epithelial tissue that develops to the adult wing blade, wing hinge and part of the dorsal body wall. During larval development, epithelial cells in the wing disc undergo rapid proliferation, leading to enlargement of the disc size(4, 5). Perturbation in cell proliferation results in adult wings with abnormal size. In addition to proliferation, cells across different regions of the wing disc gradually acquire distinct identities, establishing the anterior-posterior, dorsal-ventral, proximal-distal axes. Beginning in the mid-third instar, cells in the wing pouch, the region that gives rise to the adult wing blade, commit to either pro-vein and intervein cell fates. These populations later form the vein and intervein regions of the adult wing blade, respectively(5, 6). Owing to its well-characterized development and genetic tools, the *Drosophila* wing disc has served as a powerful model system for studying the molecular mechanisms regulating tissue growth and patterning.

*Drosophila* Ciz1 (encoded by *CG8108*) shares approximately 30% homology with mammalian CDKN1A-interacting zinc finger protein 1 (CIZ1). Mammalian CIZ1 is a nuclear matrix protein that participates in the assembly of the DNA pre-replication complex, DNA replication initiation and G1/S transition(7–9). Knockdown of *CIZ1* results in reduced entry to S phase and increased apoptosis in a couple of non-maligant and malignant cell types(7, 10, 11). Additionally, mammalian CIZ1 is a regulator of X chromosome inactivation. It interacts with the long noncoding RNA X-inactive specific transcript (Xist) to facilitate its localization to the inactive X chromosomal compartment and gene silencing(12–16). *Drosophila* Ciz1 has been detected in the nuclear matrix proteome from embryos, suggesting that like its mammalian counterpart, *Drosophila* Ciz1 is also a nuclear matrix protein(17, 18). Previous work from our group showed that knocking down *Ciz1* in *Drosophila* wing disc resulted in elevated apoptosis and reduced survival after executioner caspase activation(19). However, the role of *Drosophila* Ciz1 in organ development is not clear.

Here, we demonstrate that Ciz1 is an essential regulator of wing development. Knocking down *Ciz1* in wing discs results in reduced wing size and abnormal vein patterning. Mechanistic studies reveal that reduction in Ciz1 expression causes accumulation of donut-shaped mitochondria and elevated oxidative stress, which activates JNK signaling to cause G2 arrest and enhances EGFR signaling to induce extra vein formation.

## Results

### Knocking down *Ciz1* results in small wings with thickened and ectopic veins

To investigate the role of Ciz1 in development, we first evaluated the phenotype of transheterozygous *Ciz1^KG05452^/Ciz1^EY14316^* mutants (referred to as *Ciz1^−/−^*). *Ciz1^−/−^*animals hatched normally but exhibited growth retardation and larval lethality (Supplementary figure S1A & B). We then generated a genomic rescue construct Ciz1-GR carrying the genomic region from ∼1kb upstream of the 5’ UTR of Ciz1 to ∼1kb downstream of the 3’ UTR of Ciz1, and inserted it into attP40 site. Expression of Ciz1-GR rescued the lethality of *Ciz1^−/−^* flies (Supplementary figure S1C). The morphology of the rescued flies was comparable to that of *w^1118^* controls (Suppplementary figure S1D-F). These data suggest that Ciz1 is critical for *Drosophila* development.

Because *Ciz1* mutation caused larval lethality, to investigate the role of Ciz1 in *Drosophila* wing development, we used *nub-Gal4*, which is expressed in the pouch of the wing disc, to drive the expression of *Ciz1 RNAi* (BDSC# 27562). Ciz1 is ubiquitously expressing in the whole wing disc (Supplementary figure S2A). Expressing *Ciz1 RNAi* with *nub-Gal4* strongly reduced Ciz1 level in the pouch (Supplementary figure S2B) and resulted in dramatic reduction in adult wing size (Figure 1A & B). To determine whether the reduced wing size is due to decreased cell number or cell size, we quantified the average cell size and cell number in wings. The total cell number strongly decreased while the average cell size increased in *Ciz1* knockdown wings (Figure 1C & D). In addition to the small size, *Ciz1* knockdown wings exhibited vein defects, including significantly increased vein thickness (Figure 1A & E) and formation of ectopic vein materials, especially in the region between L2 and L3 (Figure 1A & F). Expression of *Ciz1* together with *Ciz1 RNAi* rescued all the wing defects (Figure 1A-F, Supplementary figure S1D), suggesting that the wing defects are due to loss of Ciz1. We also employed another RNAi (VDRC35344, designated as *Ciz1 RNAi.2*) that targets a different region in *Ciz1* mRNA. *Ciz1 RNAi.2* displayed stronger knockdown efficiency than *Ciz1 RNAi* (Supplementary figure S2C & E). When expressed in the pouch, it caused stronger reduction in wing size and more severe vein defects (Supplementary figure S3). These data together suggest that Ciz1 is an essential regulator of wing development.

**Figure 1.**
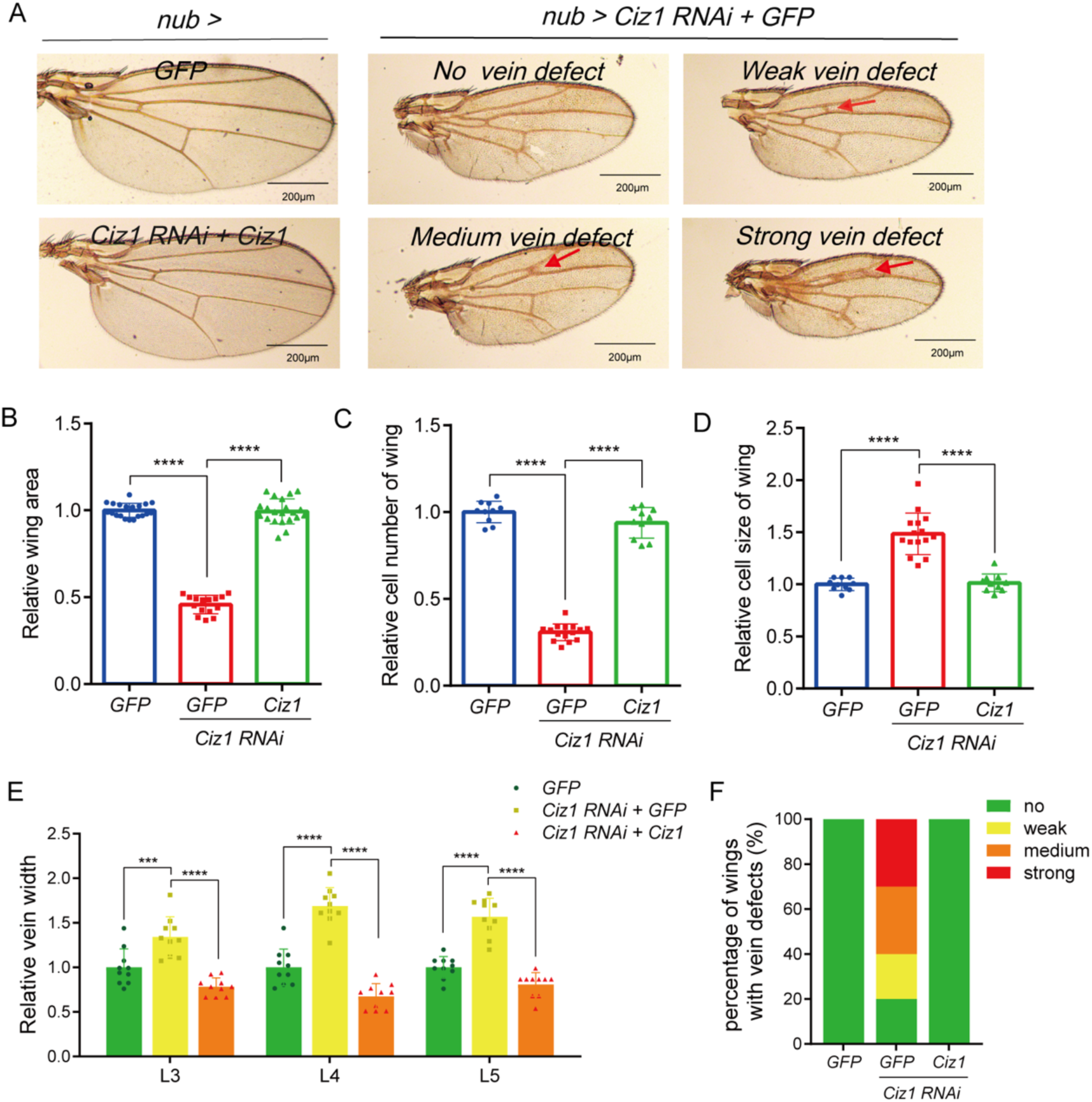
Ciz1 is essential for wing development. (A) Representative images of wings from *nub-Gal4 UAS-GFP* (*nub> GFP*), *nub-Gal4 UAS-Ciz1 RNAi UAS-GFP* (*nub> Ciz1 RNAi+GFP*) and *nub-Gal4 UAS-Ciz1 RNAi UAS-Ciz1* (*nub> Ciz1 RNAi+Ciz1*) flies. Red arrows point to ectopic veins between L2 and L3. Scale bar, 200 μm. (B-D) Quantification of wing size (B), cell number (C) and average cell size (D) in *nub> GFP*, *nub> Ciz1 RNAi+GFP* and *nub> Ciz1 RNAi+Ciz1* adult wings. n=20, 15, 21 in B. n=10, 15, 10 in C. n=10, 15, 10 in D. (E) Quantification of the width of L3, L4 and L5 in *nub> GFP* (n=10), *nub> Ciz1 RNAi+GFP* (n=10) and *nub> Ciz1 RNAi+Ciz1* (n=10) adult wings. (F) Quantification of the percentage of wings with ectopic veins. The representative images of different categories are shown in (A). n= 20 (*nub> GFP*), 40 (*nub> Ciz1 RNAi + GFP*), 32 (*nub> Ciz1 RNAi + Ciz1*). **: *P*< 0.001; ****: *P*< 0.0001. The detailed genotypes of samples in this figure are listed in Supplementary table S4.

### Knockdown of *Ciz1* induces G2 arrest

Previously we found that knocking down *Ciz1* induced apoptosis in wing discs(19) (Supplementary figure S4A). To determine whether the reduced wing size caused by *Ciz1* knockdown is due to increased apoptosis, we suppressed apoptosis in *Ciz1* knockdown discs by overexpressing the apoptosis inhibitor protein Diap1. Overexpression of Diap1 completely blocked apoptosis in *Ciz1* knockdown tissues (Supplementary figure S4A) but did not rescue the reduced wing size (Supplementary figure S4B & C), suggesting that the smaller wings are not due to increased apoptosis.

We then checked if knocking down *Ciz1* affects proliferation. Phosphor-histone 3 (pH3) is a commonly-used marker for mitosis. Knocking down *Ciz1* using *en-Gal4*, which is expressed in the posterior compartment, led to decreased pH3^+^ cells. This phenotype was rescued by restoring Ciz1 level (Figure 2A-D), suggesting that Ciz1 is required for cell proliferation. Consistent with the pro-proliferation role of Ciz1, we hardly obtained *Ciz1^−/−^* clones in wing discs using FLP/FRT system, while control clones frequently showed up. Expression of Ciz1-GR recovered *Ciz1^−/−^* clones (Supplementary figure S5A-C, F), suggesting that loss of Ciz1 causes growth disadvantage.

**Figure 2.**
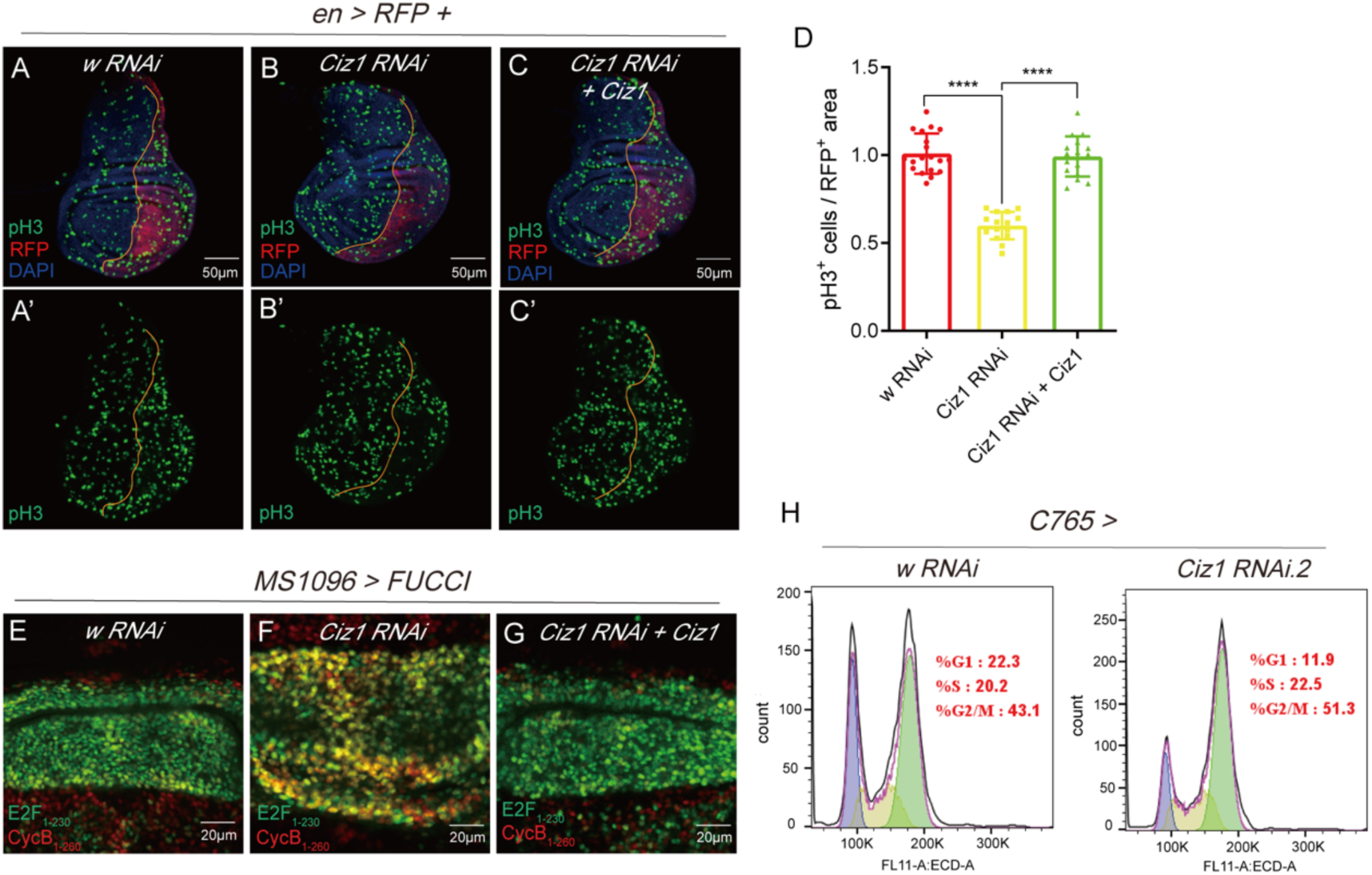
Knockdown of *Ciz1* suppresses proliferation. (A-C) The representative images of pH3 staining in wing discs. Transgenes were expressed in the posterior compartment, which is marked by RFP. The orange line marks the anterior-posterior boundary. Discs expressing RNAi targeting *white* (*w*) gene were used as control. Scale bar, 50 μm. (D) Quantification of the density of pH3^+^ cells in the posterior compartment. n= 19 (*en > w RNAi*), 15 (*en > Ciz1 RNAi*), 15 (*en > Ciz1 RNAi + Ciz1*). ****: *P*< 0.0001. (E-G) Cell cycle analysis by FUCCI reporter in wing discs. All transgenes were expressed in the wing pouch by *MS1096-Gal4*. Scale bar, 20 μm. (H) Flow cytometry analysis of DNA content to show cell cycle distribution in wing discs. The detailed genotypes of samples in this figure are listed in Supplementary table S4.

Mammalian CIZ1 is involved in DNA replication and G1/S transition(7, 20, 21). To determine whether *Drosophila* Ciz1 also regulates DNA replication, we performed EdU incorporation assay. Expression of *Ciz1 RNAi* did not affect EdU incorporation, suggesting Ciz1 is not required for DNA synthesis in *Drosophila* (Supplementary figure S6). To investigate how Ciz1 influences cell cycle, we adopted the FUCCI reporter. The FUCCI reporter marks cells in G1 and late M phase green, cells in S phase red, and cells in G2 and early M phase yellow(22). Knocking down *Ciz1* resulted in more yellow cells, which means more cells in G2 and early M phase (Figure 2E & 2F). Co-expression of Ciz1 suppressed the elevated yellow cells (Figure 2G). Flow cytometry analysis of DNA content also displayed a larger G2/M fraction in wing disc cells with *Ciz1* knocked down (Figure 2H). Considering pH3 staining showed reduced cells in M phase, the FUCCI and DNA content analysis data suggest that knocking down *Ciz1* induces G2 arrest. Thus, different from mammalian CIZ1, *Drosophila* Ciz1 is required for G2/M transition, rather than G1/S transition.

### Knockdown of *Ciz1* induces accumulation of donut-shaped mitochondria

To investigate the mechanism underlying Ciz1 regulation of wing development, we performed RNA sequencing on *Ciz1* knockdown and control discs. Knocking down *Ciz1* resulted in 86 genes upregulated and 88 genes downregulated (|Fold change| ≥ 2, *P* < 0.05, Figure 3A, Dataset S1). Interestingly, GSEA analysis revealed that the differentially expressed genes were enriched in GO terms related to mitochondrial protein complex, respiratory electron transport chain and oxidative phosphorylation (Figure 2B-E, Dataset S1). These data suggest that Ciz1 deficiency may affect mitochondria. Therefore, we examined mitochondrial morphology in wing discs with or without *Ciz1* knockdown. Using transmission electron microscope, we surprisingly found the existence of donut-shaped mitochondria in *Ciz1* knockdown discs, which were hardly captured in control discs (Figure 3F-I). To better quantify the effect of Ciz1 deficiency on mitochondrial morphology, we utilized *UAS-mito:GFP* to label mitochondria. Given that the columinar cells in wing discs are too small to view mitochondrial morphology, we expressed mito:GFP in the peripodial membrane using *C311-Gal4*. While mitochondria in control discs are elongated, *Ciz1* knockdown discs exhibited markedly increased donut-shaped mitochondria (Figure 3J-L).

**Figure 3.**
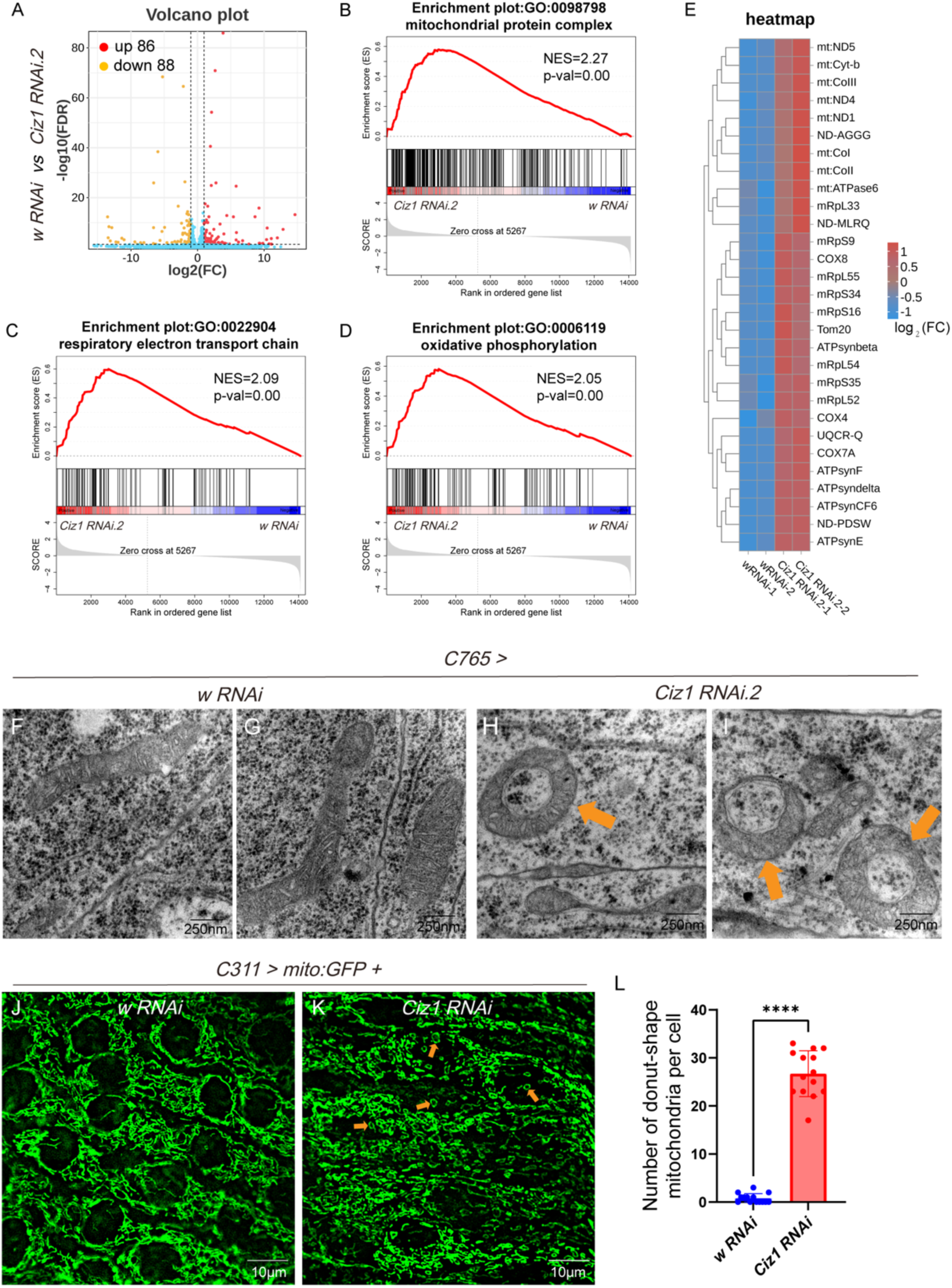
Ciz1 deficiency results in increased donut-shaped mitochondria. (A) Volcano plot showing differentially expressed genes in *Ciz1* knockdown discs compared to control discs. |Fold change| ≥ 2, *P* value < 0.05. (B-D) GSEA analysis results. (E) Heat map of expression of some mitochondria-related genes from RNAseq. (F-G) Representative TEM images. The orange arrow points to examples of donut-shaped mitochondria. Scale bar, 250nm. (J-K) Representative fluorescent images showing the morphology of mitochondria. The orange arrow points to examples of donut-shaped mitochondria. Scale bar, 10 μm. (L) Quantification of donut-shaped mitochondria in each cell. n= 14. The detailed genotypes of samples in this figure are listed in Supplementary table S4.

### Ciz1 deficiency disrupts wing development by inducing oxidative stress

Mitochondrial abnormality is frequently associated with oxidative stress(23). We used dichlorofluorescin diacetate (DCFH-DA) to detect ROS in wing discs. Knockdown of *Ciz1* markedly elevated ROS in wing discs (Figure 4A & B). Moreover, we employed GstD1:GFP, a commonly-used reporter for oxidative stress in *Drosophila*(24). We found that knocking down *Ciz1* resulted in upregulated expression of GstD1:GFP, which was rescued by expression of Ciz1 (Figure 4C-E), suggesting that reduced Ciz1 expression induces oxidative stress in wing discs.

**Figure 4.**
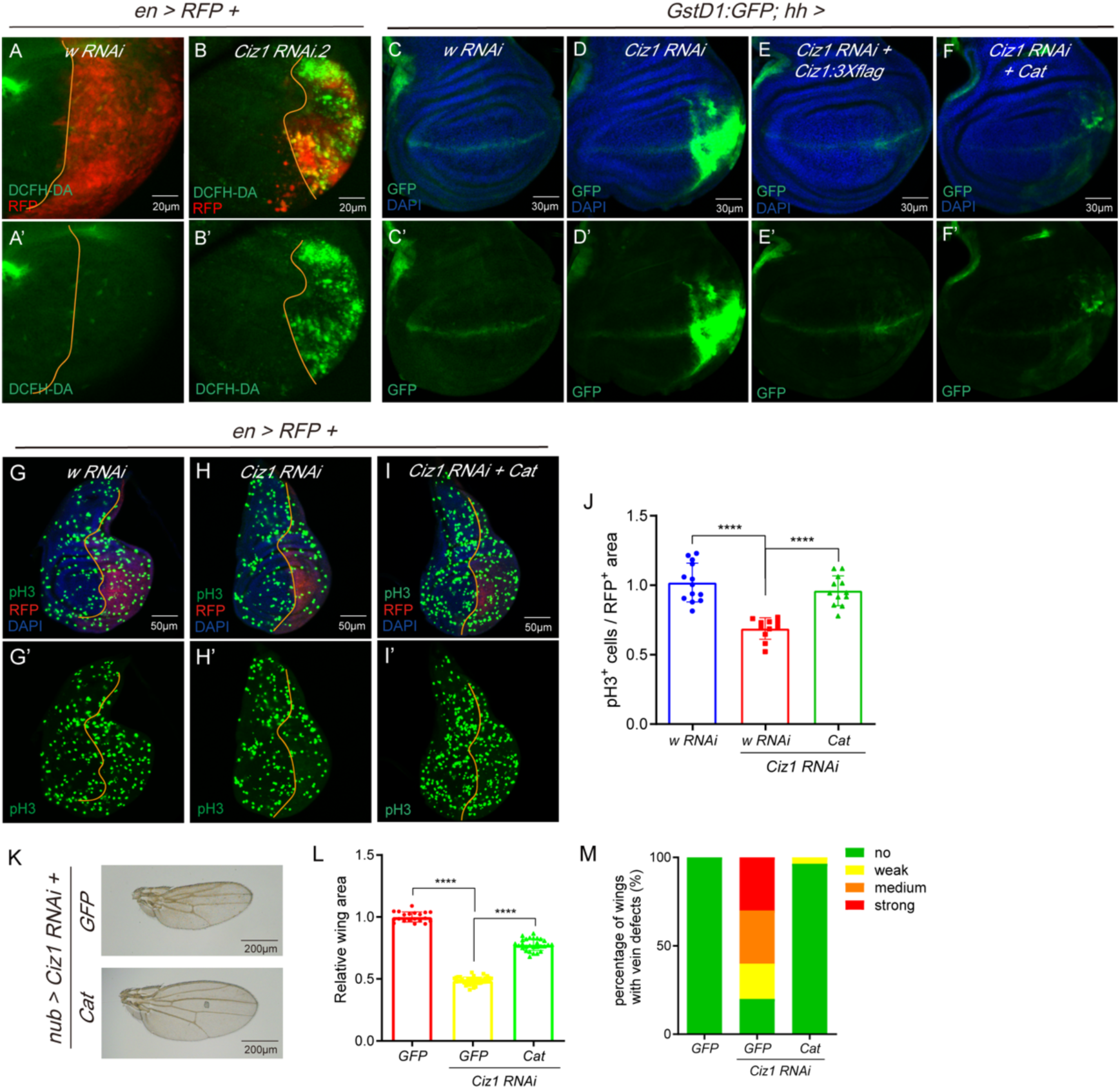
Knocking down *Ciz1* disrupts wing development by inducing oxidative stress. (A-B) The representative images of DCFH-DA staining showing ROS in wing discs with the indicated transgenes expressed in the posterior compartment labeled by RFP. Scale bar, 20 μm. (C-F) The representative images showing GstD1:GFP expression in wing discs with the indicated transgenes expressed in the posterior compartment. (G-I) The representative images of pH3 staining in wing discs with the indicated transgenes expressed in the posterior compartment marked by RFP. The orange line marks the anterior-posterior boundary. Scale bar, 50 μm. (J) The density of pH3^+^ cells in the posterior compartment. n= 13 (*en> w RNAi*), 11 (*en> Ciz1 RNAi + w RNAi*), 12 (*en> Ciz1 RNAi+ Cat*). (K) The representative images of adult wings. Scale bar, 200 μm. (L) Quantification of wing size. n= 20 (*nub> GFP*), 40 (*nub> Ciz1 RNAi + GFP*), 28 (*en> Ciz1 RNAi + Cat*). ****: *P*< 0.0001. (M) The percentage of wings with ectopic veins in different genotypes. The representative images of each category is shown in Fig 1A. The detailed genotypes of samples in this figure are listed in Supplementary table S4.

To determine whether the elevated oxidative stress is responsible for the wing defects caused by *Ciz1* knockdown, we suppressed oxidative stress by overexpressing Catalase (Cat), an enzyme involved in clearing ROS (25), in *Ciz1* knockdown discs (Figure 4F). Cat overexpression restored proliferation in *Ciz1* knockdown discs (Figure 4G-J) and significantly rescued the size defect of adult wings (from 48% of the wild-type wing size to 77.6%, Figure 4K & L). Furthermore, the vein defects caused by *Ciz1* knockdown were also rescued by overexpressing Cat (Figure 4K & M). These data together indicate that reducing Ciz1 expression disrupts wing development by inducing oxidative stress.

### Loss of Ciz1 suppresses proliferation through activating JNK

Oxidative stress often activates c-Jun NH2-terminal kinase (JNK)(26, 27) and JNK has been reported to induce G2 arrest in response to wing discs damage(28). Thus, we employed two commonly-used reporters, *puc-lacZ*(29, 30) and *TRE:DsRed*(31) to assess JNK activity upon *Ciz1* knockdown. Knocking down *Ciz1* led to elevated expression of both reporters, which was suppressed by co-expression of Ciz1 (Figure 5A-C, G-I), suggesting JNK activation in Ciz1-deficient tissues. Moreover, overexpression of Cat suppressed JNK activation in *Ciz1* knockdown tissues (Figure 5D & J), indicating that Ciz1 deficiency activates JNK through inducing oxidative stress.

**Figure 5.**
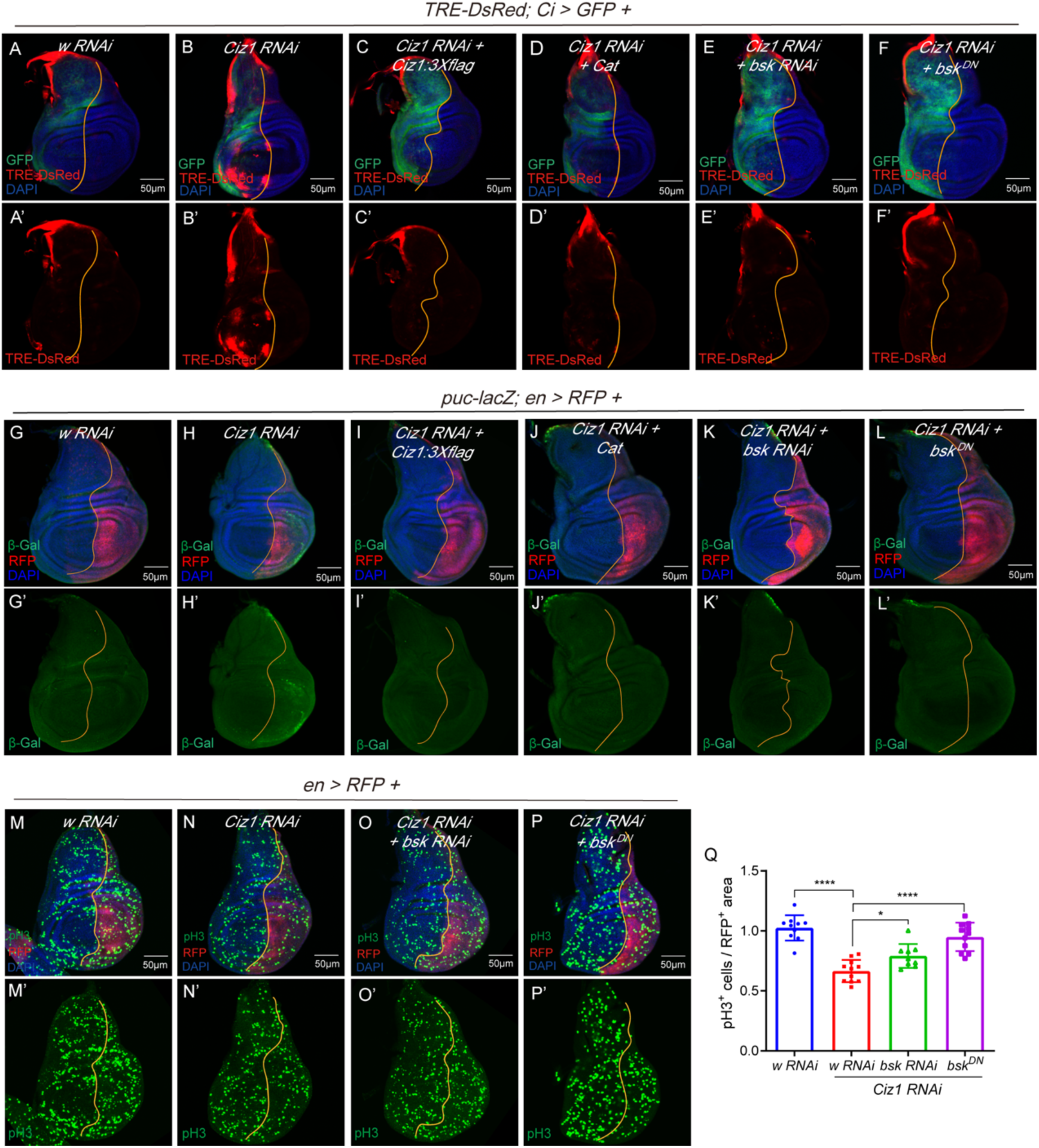
Knocking down *Ciz1* suppresses proliferation through activating JNK. (A-F) The representative images showing *TRE-DsRed* expression in wing discs with the indicated transgenes expressed in the anterior compartments labeled by GFP. The orange line marks the anterior-posterior boundary. Scale bar, 50 μm. (G-L) The representative images showing *puc-lacZ* expression in wing discs with the indicated transgenes expressed in the posterior compartment labeled by RFP. The orange line marks the anterior-posterior boundary. Scale bar, 50 μm. (M-P) The representative images of pH3 staining in wing discs with the indicated transgenes expressed in the posterior compartments labeled by RFP. The orange line marks the anterior-posterior boundary. Scale bar, 50 μm. (Q) The density of pH3^+^ cells in the posterior compartment. n= 10 (*en> w RNAi*), 11 (*en> Ciz1 RNAi + w RNAi*), 9 (*en> Ciz1 RNAi+ bsk RNAi*), 11 (*en> Ciz1 RNAi+ bsk^DN^*). *: *P*< 0.05; ****: *P*< 0.0001. The detailed genotypes of samples in this figure are listed in Supplementary table S4.

To investigate whether JNK activation mediated proliferation suppression caused by *Ciz1* knockdown, we used an RNAi against *basket* (*bsk*), the gene encoding *Drosophila* JNK, and a dominant-negative *bsk* (*bsk^DN^*) to block JNK signaling (Figure 5E, F, K, L). Expressing *bsk RNAi* or *bsk^DN^* in *Ciz1 RNAi* discs restored the percentage of pH3^+^ cells (Figure 5M-Q), indicating that Ciz1 suppresses JNK to maintain cell proliferation in wing discs.

### Knockdown of *Ciz1* induces abnormal vein formation through activating EGFR-ERK signaling

Besides reduced wing size, wings with *Ciz1* knocked down exhibited formation of ectopic vein materials, which can be rescued by clearing ROS (Figure 1F & 4M). The epidermal growth factor receptor (EGFR) - extracellular signal-regulated kinase (ERK) signaling plays a critical role in vein formation(6). Hence, we checked whether knocking down *Ciz1* affects EGFR signaling activity. In late third instar wing discs, EGFR signaling is activated in the provein region, as shown by the EGFR pathway activity reporter, *argos-lacZ* (Figure 6A). We found that knocking down *Ciz1* using *ap-Gal4* led to elevated *argos-lacZ* expression (Figure 6B), which was suppressed by expression of Ciz1 (Figure 6C). These results indicate that Ciz1 functions as an inhibitor of EGFR signaling. Overexpression of Cat suppressed the increased *argos-lacZ* expression in *Ciz1* knockdown discs (Figure 6D), suggesting that the EGFR activation relies on the elevated oxidative stress.

**Figure 6.**
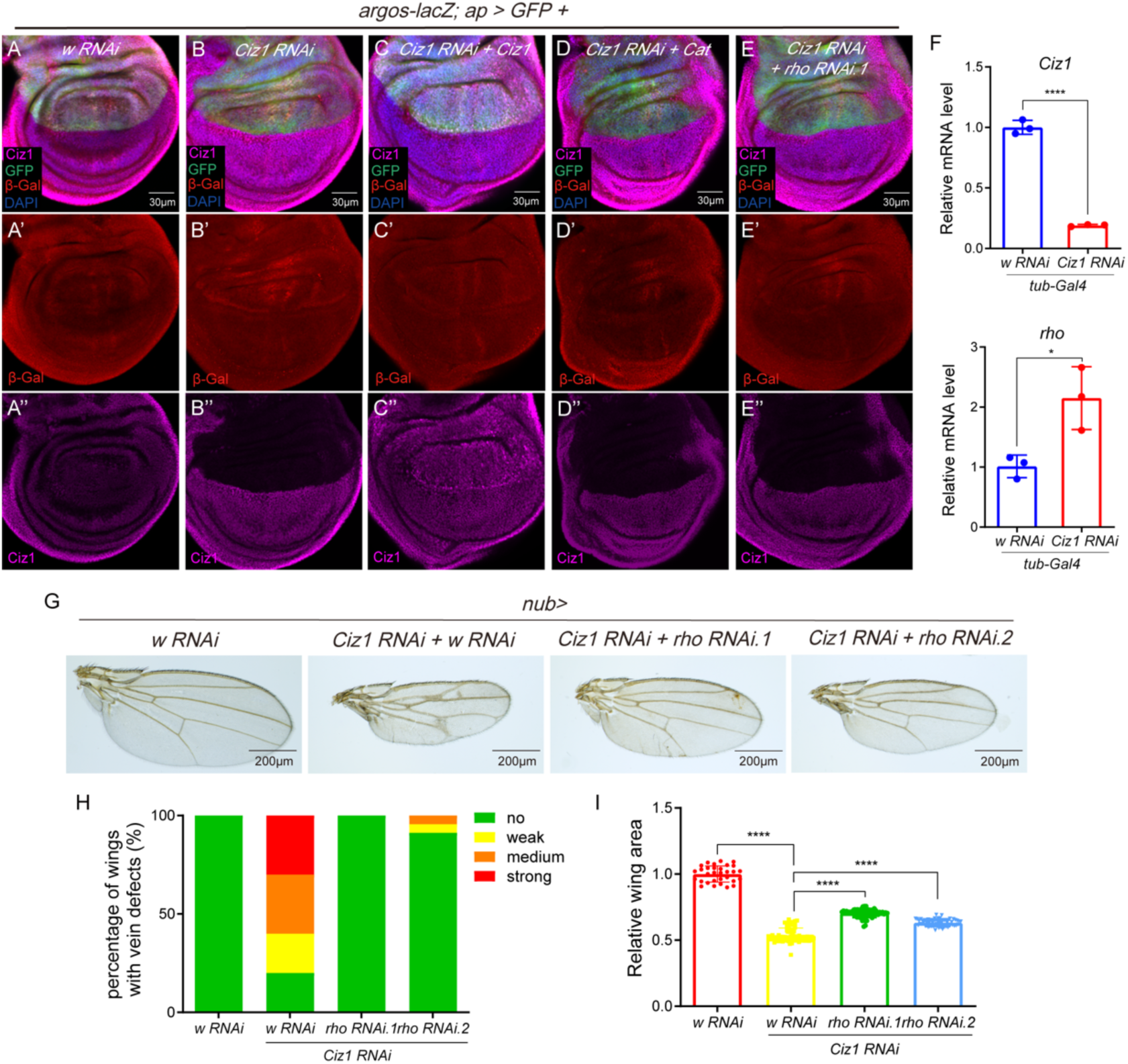
Knocking down *Ciz1* induces ectopic veins through activating EGFR-ERK signaling. (A-E) The representative images showing *argos-lacZ* expression in wing discs with the indicated transgenes expressed in the dorsal compartment labeled by GFP. Scale bar, 30 μm. (F) The mRNA levels of *rho* and *Ciz1* detected by qRT-PCR. n=3. Each biological replicate was prepared from 100 discs. (G) The representative images of adult wings. Scale bar, 200 μm. (H) The percentage of wings with ectopic veins in different genotypes. The representative images of each category is shown in Fig 1A. (I) Quantification of adult wing size. n= 31 (*nub> w RNAi*), 68 (*nub> Ciz1 RNAi + w RNAi*), 67 (*nub> Ciz1 RNAi + rho RNAi.1*), 66 (*nub> Ciz1 RNAi + rho RNAi.2*). *rho RNAi.1* and *rho RNAi.2* are two different RNAi lines targeting *rho*. *: *P*< 0.05; ****: *P*< 0.0001. The detailed genotypes of samples in this figure are listed in Supplementary table S4.

To determine whether the enhanced EGFR signaling mediates the formation of ectopic veins, we first evaluated the effect of the heterozygous mutation of *rolled* (*rl*), the *Drosophila* ERK, on the vein defects caused by *Ciz1* knockdown. The vein defects were dramatically rescued in *rl^1^/+* flies (Supplementary figure S7A & B). We further found that knocking down *Egfr* also rescued the vein defects (Supplementary figure S7D & E). These data indicate that Ciz1 suppresses vein formation by inhibiting EGFR-ERK pathway.

The results that knocking down *Egfr* suppresses vein defects caused by *Ciz1 RNAi* suggest that Ciz1 may regulate EGFR signaling at or upstream of EGFR. We checked the expression of the upstream EGFR signaling regulators and found upregulatioin of *rhomboid* (*rho*) in *Ciz1* knockdown discs (Figure 6F). Rhomboid (Rho) is an intra-membrane serine protease that can cleave and activate EGFR ligands(32–35). Reducing *rho* in *Ciz1* knockdown discs suppressed the upregulated *argos-lacZ* (Figure 6E) and vein defects (Figure 6G & H), suggesting that Ciz1 suppresses EGFR signaling and vein defects by inhibiting expression of *rho*.

The aforementioned data reveal activation of both JNK and EGFR signaling by oxidative stress in *Ciz1* knockdown discs. We then investigated the interaction between EGFR signaling and JNK signaling. Knocking down *Egfr* failed to suppress elevated JNK activity in *Ciz1* knockdown discs and similarly, reducing *bsk* expression did not affect EGFR activity (Supplementary figure S8), indicating that Ciz1 suppresses JNK pathway and EGFR signaling in parallel. Although heterozygous mutation of *rl* had no effect on wing size (Supplementary figure S7C), knocking down *Egfr* or *rho* slightly increased the size of *Ciz1* knockdown wings (Figure 6I & Supplementary figure S7F). This moderate change may be attributed to the smaller size of vein cells compared to intervein cells.

### Ciz1 relies on its N-terminal part to restrain oxidative stress and promote wing development

Analysis of Ciz1 protein sequence identified two zinc finger motifs (located at 476-498aa and 571-593aa) (Figure 7A). To investigate whether these motifs are required for Ciz1 regulation of oxidative stress and wing development, we generated a transgenic fly line carrying *UAS-Ciz1^ΔZF^*, which encodes a mutant form of Ciz1 with 476-593aa deleted (Figure 7A). Expression of Ciz1^ΔZF^ effectively rescued all the defects caused by *Ciz1* knockdown including the reduced wing size, abnormal vein patterning (Figure 7B-D), increased oxidative stress (Figure 7E & F), decreased proliferation (Supplementary figure S9A-C, G, H) and elevated JNK (Supplementary figure S10A & H) and EGFR activation (Supplementary figure S11A-D), to an extent comparable to expression of full-length Ciz1. To further confirm the dispensability of the zinc finger motifs in Ciz1, we generated a genomic rescue construct Ciz1-GR^ΔZF^, in which the region encoding 476-593aa was deleted from Ciz1-GR construct. Expression of Ciz1-GR^ΔZF^ rescued the lethality of *Ciz1^−/−^* mutants (Supplementary figure S1C-F) and recovered *Ciz1^−/−^* clones in wing discs (Supplementary figure S5D & S5F). These data together suggest that the zinc finger motifs are not required for Ciz1 regulation of wing development.

**Figure 7.**
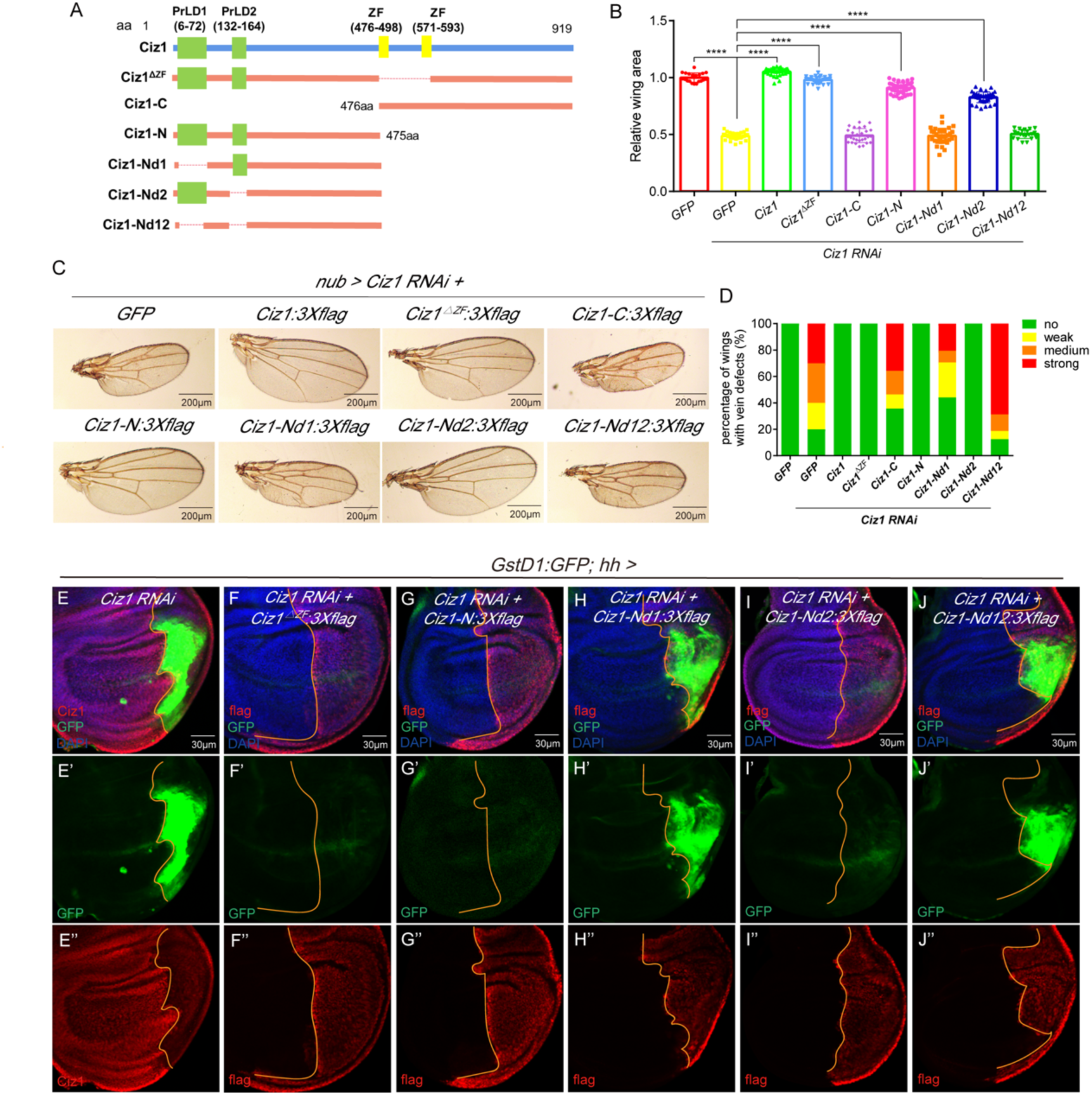
Ciz1 regulation of wing development relies on its N-terminal part. (A) The schematic showing different fragments of Ciz1. (B) Quantification of wing size. n= 20 (*nub> GFP*), 40 (*nub> Ciz1 RNAi + GFP*), 32 (*nub> Ciz1 RNAi + Ciz1*), 29 (*nub> Ciz1 RNAi + Ciz1^ΔZF^*), 28 (*nub> Ciz1 RNAi + Ciz1-C*), 39 (*nub> Ciz1 RNAi + Ciz1-N*), 35 (*nub> Ciz1 RNAi + Ciz1-Nd1*), 35 (*nub> Ciz1 RNAi + Ciz1-Nd2*), 21 (*nub> Ciz1 RNAi + Ciz1-Nd12*). ****: *P*< 0.0001. (C) The representative images of adult wings. Scale bar, 200 μm. (D) The percentage of wings with ectopic veins in different genotypes. The representative images of each category is shown in Fig 1A. n= 20 (*nub> GFP*), 40 (*nub> Ciz1 RNAi + GFP*), 32 (*nub> Ciz1 RNAi + Ciz1*), 29 (*nub> Ciz1 RNAi + Ciz1^ΔZF^*), 28 (*nub> Ciz1 RNAi + Ciz1-C*), 39 (*nub> Ciz1 RNAi + Ciz1-N*), 34 (*nub> Ciz1 RNAi + Ciz1-Nd1*), 35 (*nub> Ciz1 RNAi + Ciz1-Nd2*), 16 (*nub> Ciz1 RNAi + Ciz1-Nd12*). (E-J) The representative images showing GstD1:GFP expression in wing discs with the indicated transgenes expressed in the posterior compartment marked by either reduced Ciz1 expression (E) or expression of flag-tagged proteins (F-J). The orange line marks the anterior-posterior boundary. Scale bar, 30 μm. The detailed genotypes of samples in this figure are listed in Supplementary table S4.

To determine which part of Ciz1 is essential for its function in wing development, we generated *UAS-Ciz1-N* and *UAS-Ciz1-C* flies, which, when combined with Gal4, can express the N-terminal part (1-475aa) and the C-terminal part (476-919aa) of Ciz1 protein, respectively (Figure 7A). Expression of Ciz1-N fully rescued all the defects resulting from *Ciz1* knockdown, whereas Ciz1-C failed (Figure 7B-D, G, Supplementary figure S9-S11), indicating that the N-terminal half is necessary and sufficient for Ciz1 function in wing development.

The N-terminal region contains two prion-like domains (PrLDs, 6-72aa and 132-164aa). To assess their functional relevance, we generated transgenic flies expressing three deletion variants of the N-terminal region, *UAS-Ciz1-Nd1* (deleting 2-72aa), *UAS-Ciz1-Nd2* (deleting 132-164aa) and *UAS-Ciz1-Nd12* (deleting both regions) (Figure 7A). Expression of Ciz1-Nd2 rescued the size and vein defects in adult wings caused by *Ciz1* knockdown, whereas Ciz1-Nd1 and Ciz1-Nd12 did not (Figure 7B-D). Consistently, Ciz1-Nd2 but not Ciz1-Nd1 or Ciz1-Nd12 suppressed oxidative stress and restored proliferation and cell cycle progression in *Ciz1* knockdown discs (Figure 7H-J, Supplementary figure S9-S11). These data indicate that the first PrLD is essential for Ciz1 function in wing development.

### Ciz1 functions in cytosol to restrain oxidative stress

Previous proteomic studies identified Ciz1 in nuclear proteins(17, 18). Immunostaining of both endogenous and exogenous Ciz1 revealed that it predominantly located in the nucleus (Figure 8A & B). We further found that Ciz1-N but not Ciz1-C is nuclear localized (Figure 8C & D), suggesting that the nuclear localization signal (NLS) resides in the N-terminal part. By expressing different fragments of Ciz1-N in cultured cells, we figured out that NLS of Ciz1 resided in 73-131aa, between the two PrLDs (Supplementary figure S12). This was further confirmed by expressing Ciz1 with 73-131aa deleted (Ciz1^ΔNLS^) and Ciz1-NLSC, which fused the 73-131aa to the C-terminal half, in the wing disc. Ciz1^ΔNLS^ lost the nuclear localization while fusion of 73-131aa brought the C-terminal half into the nucleus (Figure 8E & F). To determine whether Ciz1 functions in the nucleus to regulate wing development, we expressed Ciz1^ΔNLS^ in the *Ciz1* knockdown disc. Surprisingly, expressing Ciz1^ΔNLS^ suppressed the elevated oxidative stress, JNK activation and EGFR activation in *Ciz1* knockdown tissues (Figure 8G & H, Supplementary figure S10, S11). Ciz1^ΔNLS^ also significantly increased the density of pH3^+^ cells in *Ciz1*-deficient compartments (Figure 8I-K). Consistently, we generated transgenic flies carry genomic rescue construct with the region encoding NLS deleted (Ciz1-GR-ΔNLS). Although Ciz1-GR-ΔNLS failed to rescue the lethality of *Ciz1^−/−^* flies (Supplementary figure S1C), it rescued the growth of *Ciz1^−/−^*clones in wing discs (Supplementary figure S5E & F), suggesting that although the nuclear localization of Ciz1 is required for larval survival, the cytosolic Ciz1 is sufficient to maintain cell proliferation in wing discs. Furthermore, expressing Ciz1^ΔNLS^ in *Ciz1* knockdown discs almost completely rescued the vein defects and restored the wing size from 47.6% of the wild-type to 71.8%, similar to what achieved by inhibition of oxidative stress (Figure 8L-N). Taken together, Ciz1 mainly functions in cytosol to prevent oxidative stress and maintain wing development.

**Figure 8.**
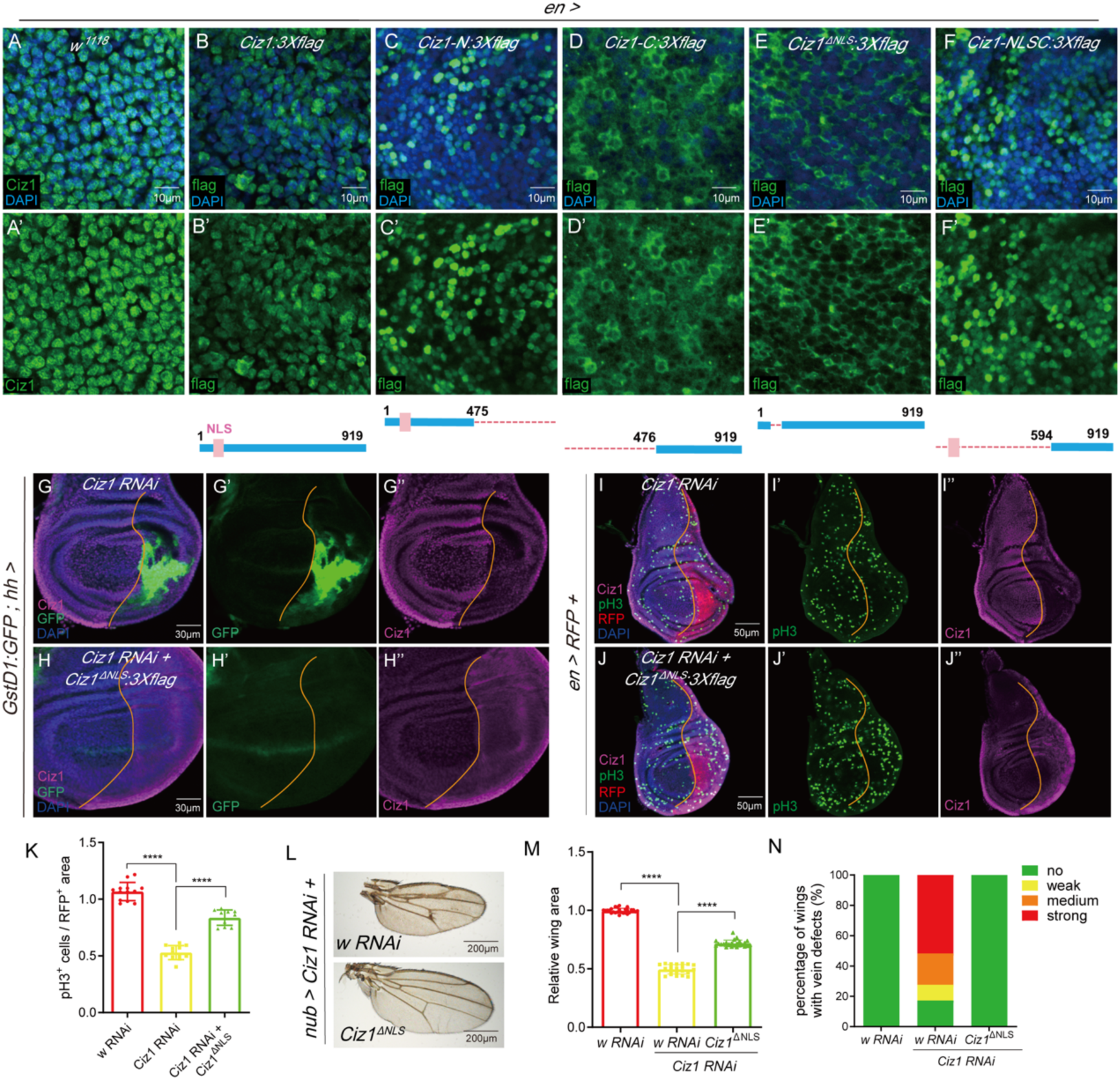
Ciz1 mainly functions in the cytosol to restrain oxidative stress. (A-F) The representative images showing the subcellular localization of endogenous Ciz1 or exogenous flag-tagged Ciz1 or Ciz1 fragments. Scale bar, 10 μm. The schematics below (B-F) show the composition of the fragments. (G-H) Representative images showing GstD1:GFP expression in wing discs with the indicated transgenes expressed in the posterior compartments. Scale bar, 30 μm. (I-J) Representative images of pH3 staining in wing discs with the indicated transgenes expressed in the posterior compartments marked by RFP. Scale bar, 30 μm. The orange lines mark the anterior-posterior boundary. (K) Quantification of the density of pH3^+^ cells in the posterior compartments. n= 13 (*en> w RNAi*), 12 (*en> Ciz1 RNAi*), 11 (*en> Ciz1 RNAi + Ciz1^ΔNLS^*). (L) Representative images of adult wings. Scale bar, 200 μm. (M) Quantification of adult wing size. n= 26 (*nub> w RNAi*), 21 (*nub> Ciz1 RNAi + w RNAi*), 23 (*nub> Ciz1 RNAi + Ciz1^ΔNLS^*). (N) The percentage of wings with ectopic veins in different genotypes. The representative images of each category is shown in Fig 1A. n= 26 (*nub> w RNAi*), 29 (*nub> Ciz1 RNAi + w RNAi*), 23 (*nub> Ciz1 RNAi + Ciz1^ΔNLS^*). ****: *P*< 0.0001. The detailed genotypes of samples in this figure are listed in Supplementary table S4.

## Discussion

In this study, we identify *Drosophila* Ciz1 as a crucial regulator that prevents oxidative stress, thereby ensuring proper wing development. Loss of Ciz1 leads to accumulation of donut-shaped mitochondria and increased oxidative stress in the wing disc epithelium, which in turn triggers activation of JNK and EGFR signalings, leading to severely reduced wing size and formation of ectopic veins. By generating a series of Ciz1 fragments, we demonstrate that although Ciz1 is predominantly located in the nucleus, it suppresses oxidative stress in the cytosol (Figure 9).

**Figure 9.**
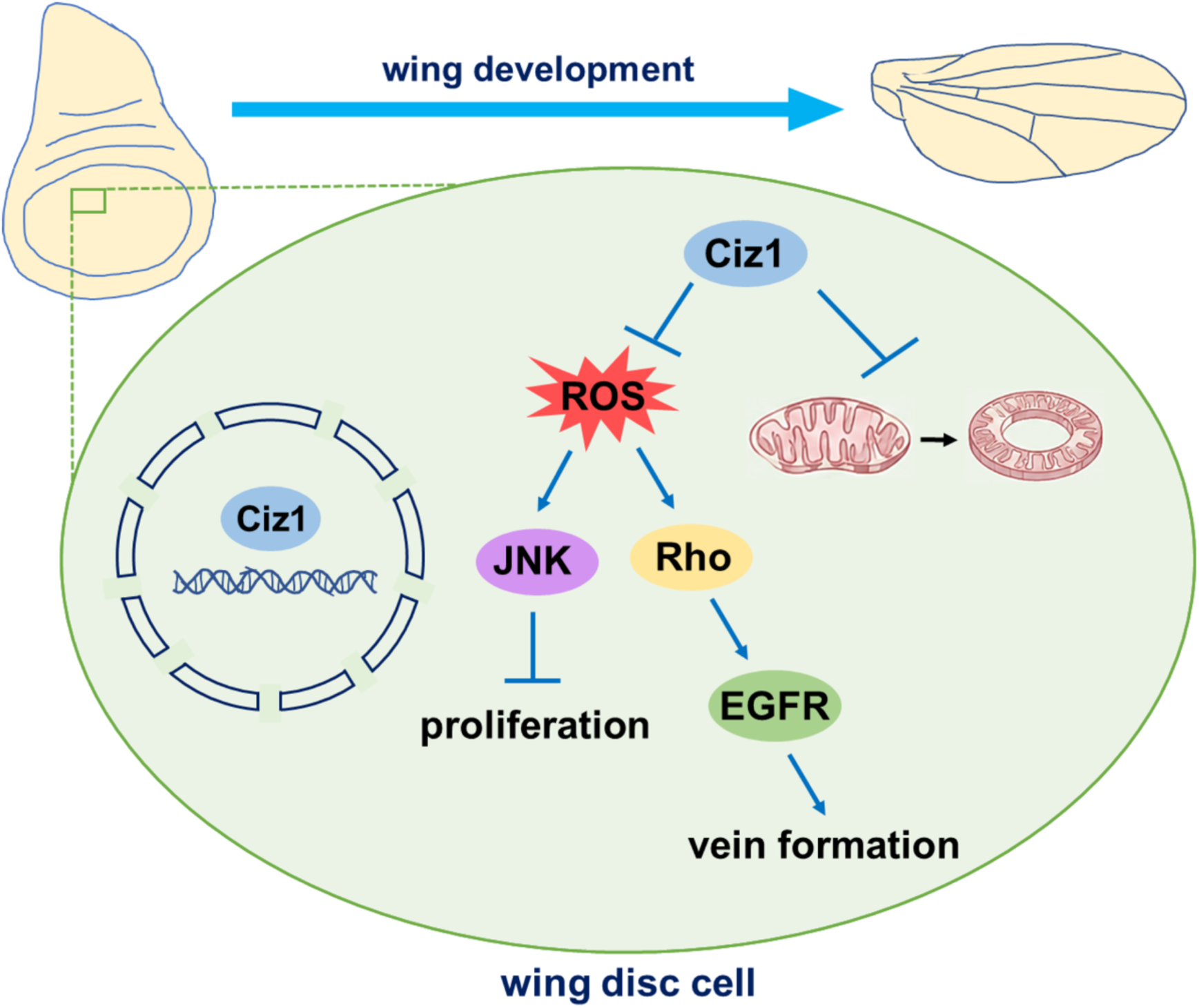
The mechanisms by which Ciz1 regulates wing development.

Our work show that knockdown of *Ciz1* results in an approximately 50% reduction in wing size. Since organ size is determined by both the number of cells in the organ and the size of each cell(4, 36, 37), we further examined these parameters and found that *Ciz1* deficiency led to a dramatic decrease in total cell number in the wing, coupled with an increase in average cell size. The increased cell size may represent a compensatory response to the severe reduction in cell number. Although loss of Ciz1 affected both cell death and proliferation, we demonstrated that blocking cell death did not rescue the wing size defect, suggesting that Ciz1 controls organ size primarily through regulation of proliferation. In mouse heart, cardiomyocytes exhibit lower CIZ1 level compared to the non-cardiomyocytes. Overexpression of CIZ1 specifically in cardiomyocytes leads to enlarged hearts with increased cardiomyocyte number and reduced cardiomyocyte size(38). This work, together with ours, suggests a conserved role of CIZ1 in organ size control, mediated through its regulation of proliferation.

The role of mammalian CIZ1 in cell proliferation has been studied by several groups. CIZ1 was first identified in a yeast two-hybrid screen as a binding partner of p21, an important cell cycle regulator(39). Subsequent studies have demonstrated that mammalian CIZ1 can trigger DNA replication both in the cell-free system and in intact cells(7–9). However, we found that knocking down *Ciz1* in *Drosophila* wing disc did not impair DNA synthesis. Rather, it caused G2 arrest through mild activation of JNK signaling. This finding suggests that although both mammalian CIZ1 and *Drosophila* Ciz1 are important proliferation regulators, they operate through distinct molecular mechanisms.

JNK signaling is a stress-activated pathway involved in regulation of cell proliferation, apoptosis and migration. Strong JNK activation triggers apoptosis by inducing expression of pro-apoptotic genes like *hid*(40, 41). However, when apoptosis is blocked, activation of JNK displays pro-proliferative effects and drives tissue overgrowth(41, 42). JNK is also required for driving compensatory proliferation during regeneration(43–47) and for sustaining proliferation in several tumor models(42, 48–54). Recent studies indicate that mild JNK activation can lead to cell cycle arrest and proliferation suppression. Kucinski et al. demonstrated that chronic JNK activation cooperates with JAK-STAT signaling and oxidative stress to suppress proliferation of wing disc cells, thereby establishing the “loser” cell status(55). Cosolo et al. reported that JNK activation following wing disc injury induces either transient G2 stalling or a prolonged but reversible G2 arrest, which protect cells from apoptosis(28). Suppression of G2/M transition by JNK has also been reported in mammalian cells(56, 57). Our finding that JNK is responsible for proliferation suppression and G2 arrest caused by *Ciz1* knockdown further support the model that mild JNK activation can induce G2 arrest.

In addition to reduced wing size, *Ciz1* knockdown leads to formation of ectopic veins. We further demonstrate that Ciz1 prevents excessive EGFR signaling activation to ensure proper vein patterning during wing development. EGFR signaling is a key regulator of longitudinal vein formation. Upon ligand binding, EGFR activates Ras-Raf-MEK cascade, resulting in ERK activation, which instructs epithelial cells to adopt the provein fate. Numerous molecules affecting expression or activity of components within EGFR signaling have been reported to influence vein formation(6, 32, 33, 58–62). In this study, we show that knocking down *Ciz1* induces EGFR signaling activation and ectopic vein formation through upregulation of Rhomboid, an essential molecule in activation of EGFR ligands(32–35).

We further demonstrate that *Ciz1* knockdown leads to elevated ROS and oxidative stress. Scavenging ROS by overexpression Cat almost completely suppressed the formation of ectopic veins and restored wing size from ∼48.7% of the normal size to 77.6%, suggesting that oxidative stress is the primary mediator of abnormal wing development caused by Ciz1 deficiency. Oxidative stress is known to regulate multiple signaling pathways, including JNK, p38 MAP kinase and EGFR-ERK signaling(63, 64). Our study shows that inhibition of oxidative stress suppresses activation of JNK and EGFR induced by *Ciz1* knockdown, placing oxidative stress upstream of these two pathways. Yet, how oxidative stress induces JNK and EGFR activation in *Ciz1* knockdown discs needs further investigation.

Interestingly, we observed an increase in donut-shaped mitochondria in cells with reduced Ciz1 expression. Donut-shaped mitochondria have been reported in cultured cells exposed to hypoxia-reoxygenation(65), osteoblasts(66), glycolytic muscle fibers(67), aging oocytes(68) and aging neurons(69). The function of these mitochondria remains unclear, but they may represent an adaptive response to stress(70). Ahmad et al. have reported that oxidative stress can trigger the formation of donut-shaped mitochondria(23). Therefore, the donut-shape mitochondria observed in *Ciz1* knockdown cells may result from increased oxidative stress. Further efforts are required to elucidate the exact role of these morphological changes of mitochondria in wing disc development.

Ciz1 is a zinc finger protein that primarily localizes to the nucleus. However, we found that expression of Ciz1 with the two zinc finger motifs deleted in Ciz1-deficient flies can rescue all the observed developmental defects. Furthermore, although Ciz1 with a deleted nuclear localization signal cannot rescue the lethality of *Ciz1^−/−^*animals, it can suppress oxidative stress in *Ciz1* knockdown wing discs and rescued wing defects to an extent similar to that achieved by inhibition of oxidative stress. These data indicate that Ciz1 functions in the cytoplasm to regulate oxidative stress. We also found that expression of the N-terminal fragment containing the first 475 residues was sufficient to rescue all wing defects and suppressed oxidative stress caused by *Ciz1* knockdown. The N-terminal part contains two PrLDs. Deletion of the PrLD located at 6-72aa abolished the ability of the N-terminal fragment to rescue defects in *Ciz1* knockdown wings, underscoring the necessity of this region in wing development regulation. PrLDs are low-complexity domains with amino acid composition similar to yeast prion domains, which enables formation of self-templating conformers. They are present in many DNA or RNA-binding proteins and play key roles in mediating phase separation and protein aggregation(71, 72). The dependence on the PrLD and the dispensiblity of the NLS for Ciz1 function in wing development suggests that Ciz1 may suppress oxidative stress and sustain normal wing development through protein-protein or protein-nuclei acid interactions in the cytosol. Further study is necessary to elucidate the precise molecular mechanisms by which Ciz1 regulates oxdative stress and wing development.

## Materials and methods

### Fly stocks

Fly stocks used in this study were listed in Supplementary table S1. For transgenic flies made in this study, plasmids with attB sequence were injected into attP40 site by Fungene Biotechnology (Qidong, China) and UniHuaii (Zhuhai, China). All crosses were raised at 25 °C till the mid to late third instar for wing disc collection or till the adulthood for adult wing analyses. To generate clones in wing discs, larvae were heat-shocked at 37 °C for 60 minutes on day 2 after egg laying.

### Plasmids

To make plasmids expressing different flag-tagged Ciz1 fragments in flies or mammalian cells, the corresponding fragment of Ciz1 was cloned using *pJFRC7-20×UAS-Ciz1:3×flag* plasmid(19) as the template and inserted into the *pJFRC7-20×UAS* vector (made from Addgene#26220, to make transgenic flies) or the *pCDNA3.1-V5:HisB* vector (ThermoFisher Scientific, Cat# V81020, for expression in mammalian cells). To make the *Ciz1-GR* construct, the genomic region from ∼1kb upstream of the 5’ UTR of *Ciz1* to ∼1kb downstream of the 3’ UTR of Ciz1 was cloned using the genomic DNA of *w^1118^*flies as the template, and inserted into pattB vector. The *Ciz1-GR^ΔZF^* and *Ciz1-GR^ΔNLS^* constructs were then made using *Ciz1-GR* construct as the template. The primers used in PCR are listed in Supplementary table S2.

### Cell culture and transfection

HEK-293T cells were cultured in DMEM medium (Gibco, Cat# C11995500BT) with 10% fetal bovine serum (HAKATA, Cat# HB-FBS-500) in a humidified 37 °C incubator with 5% CO_2_. One day before transfection, cells were seeded in a six-well plate at the density of about 3×10^5^ cells per well. Plasmids and Lipofectamine 2000 (Thermo Fisher Scientific, Cat# 11668019) were diluted in Opti-MEM (Gibco, Cat# 31985070) separately, and incubated at room temperature (RT) for 5 min. Plasmid solution and Lipofectamine solution were then mixed and incubated at RT for 25 min. The culture medium of the cells were then replaced with the mixture of plasmids and Lipofectamine 2000. After 6-to-8-hour incubation at 37 °C, the mixture was replaced with fresh culture medium. Cells were collected for immunostaining after an additional 24-hour culture.

### Immunostaining and imaging

HEK-293T cells were seeded on coverslips until they adhered. The anterior parts of third instar larvae of the desired genotypes were removed and inverted in PBS. Samples were fixed with 4% paraformaldehyde (Sangon, Cat# E672002) for 20 minutes at RT followed by washing with PBS containing 0.3% Triton X-100 (Sangon, Cat# A110694) (PBT) for 20 minutes. The samples were then blocked with 5% goat serum (Beyotime, Cat# C0265) in PBT for 30 minutes at RT followed by overnight incubation with primary antibody solution on a shaker at 4 °C. After being washed with PBT, the samples were incubated with secondary antibodies and 5 μg/ml Hoechst (Beyotime, Cat# C1022) on a shaker at RT for 1.5 hours followed by additional wash with PBT to remove residual staining solution. The wing discs were then dissected in PBT and mounted on slides with 20 μl Fluoroshield (Sigma, Cat# F6182). For EdU / DCFH-DA assays, the samples were incubated with 10 μM EdU (Beyotime, Cat# C0071S) on a shaker at RT for 10 minutes or 10 μM DCFH-DA (Beyotime, Cat# S0033S) for 30 minutes before fixation. Images were taken using Zeiss LSM 880 confocal microscope and Zeiss LSM 980 confocal microscope. The software used in image capture is Zen Black (Zeiss). The super-resolution image was taken with HIS-SIM from CSR Biotech. The image analyses were done using ImageJ. The polyclonal antibody against *Drosophila* Ciz1 was generated by immuning rabbits with the peptide containing 530-696aa of Ciz1 by Abclonal Technology (Wuhan, China). All primary and secondary antibodies used in the immunostaining are listed in Supplementary table S3.

### Wing mounting and measurement

Adult flies with the desired genotypes were collected and stored at −20°C. For dissection, the frozen flies were placed in isopropanol at RT. The wings were carefully taken off and mounted on the slides with 20 μl of 80% glycerol. The images were taken using a Zeiss microscope. The wing size, cell size and cell number were quantified using ImageJ.

### RNA extraction, reverse transcription, and qPCR

About 50 late third instar wing discs were dissected in PBS, and immediately transferred to TRIZOL (Thermo Fisher Scientific, Cat# 15596018). The total RNA was extracted using standard TRIZOL RNA extraction protocol. Removal of genomic DNA contamination and reverse transcription were done using HiScript III RT SuperMix for qPCR (Vazyme Cat# R323). qPCR was done using ChamQ SYBR Color qPCR Master Mix (Vazyme Cat# Q411) on Bio-rad CFX96 real-time PCR detection system. The data were collected using Bio-rad CFX Manager software (Bio-rad). The primers used for qPCR were listed in Supplementary table S2.

### Flow cytometry

200 wing discs were dissected and digested with 200 μL of 9× trypsin at 37°C for 15 minutes. The digestion was terminated by adding 500 μL of culture medium. The samples were then centrifuged at 830 rpm for 3 minutes and the supernatant was discarded. The cells were resuspended with 500 μL of pre-cooled 70% ethanol and fixed at 4°C overnight. The cells were then washed, pelleted and resuspended with 300 μL PBS containing 10 μL ribonuclease and 100 μL PI staining solution (Beyotime, Cat# ST511). After 30-minute incubation, cells were filtered through a strainer and applied to flow cytometry (Beckman, CytoFLEX S). Twenty thousand cells were collected at low speed. The data were processed with FlowJo and fit to the cell cycle distribution plot.

### Transmission electron microscopy (TEM)

The anterior parts of larvae were tore off and fixed with glutaraldehyde and osmium acid. After dehydration through an ethanol gradient, resin infiltration was performed. The wing discs were then dissected out and embedded. After polymerization, samples were cut to sections of 50-70 nm and stained with uranyl acetate and lead citrate. Sections were then placed onto copper grids and flash-frozen with liquid nitrogen. All reagents and consumables used were obtained from SPI supplies (West Chester, PA, USA) and EMCN (Beijing, China). Images of mitochondria were captured using JEM-1200EX at 4000× magnification.

### RNA sequencing

For each sample, about 500 late third instar wing discs were dissected in PBS, transferred to Trizol (Thermo Fisher Scientific, Cat# 15596018) in RNase-free tubes, and store at −80°C. RNA sequencing was performed by Genedenovo Company (Guangzhou, China) and data analyses were performed using Omicsmart (Genedenovo Company).

### Statistics and reproducibility

The data were presented as mean ± standard deviation (SD). All the statistical analyses were done using GraphPad Prism 10 software. Statistical significance was determined using unpaired two-tailed t-test for two-sample comparison or one-way ANOVA for multiple-sample comparison. The Tukey test was used to derive adjusted *P*-value for multiple comparisons. *P*< 0.05 was considered statistically significant. All experiments were independently repeated at least three times to ensure reproducibility, and the representative images are shown in the figures. The sample size was listed in the figure legends.

## Data availability statement

The raw data of RNA sequencing can be accessed at NCBI Bioproject PRJNA1476231. All the other raw data supporting the findings of this study are available from the corresponding author upon request.

## Acknowledgement

We thank Dr. Zizhang Zhou, Dr. Junzheng Zhang, Dr. Xuan Guo, TsingHua Fly Center, National Drosophila Resource Center of China, Bloomington Drosophila Stock Center and Vienna Drosophila Resource Center for providing fly stocks. We thank Translational Medicine Core Facility of Shandong University and the School of Basic Medical Sciences Core Facility of Shandong University for technical support. This work was supported by National Natural Science Foundation of China (No. 32270869), Taishan Scholar Foundation of Shandong Province (tsqn202312025) and Natural Science Foundation of Shandong Province (ZR2025QB06) to G.S.

## CRediT authorship contribution statement

Xiaojiao Li: Writing - Original draft, Investigation, Data curation. Cuiping Wang: Investivation, Data curation. Yuanyuan Zhang: Validation, Formal analysis. Huan Liu: Validation, Formal analysis. Min Hou: Resources, Validation. Xiuxiu Liu: Resources, Validation. Yuanlin Su: Validation. Yanqin Gong: Resources. Huiyuan Ding: Resources. Molin Wang: Resources. Yongxin Zou: Methodology. Qiao Liu: Visualization. Yaoqin Gong: Supervision, Writing – Review & Editing. Gongping Sun: Conceptualization, Supervision, Writing – Original draft, Writing – Review & Editing.

