## Supplementary figures and tables for "Ciz1 safeguards *Drosophila* wing development by suppressing oxidative stress"

Li et al.

The file contains the following contents.

Supplementary figure S1-S12

Supplementary table S1-S4

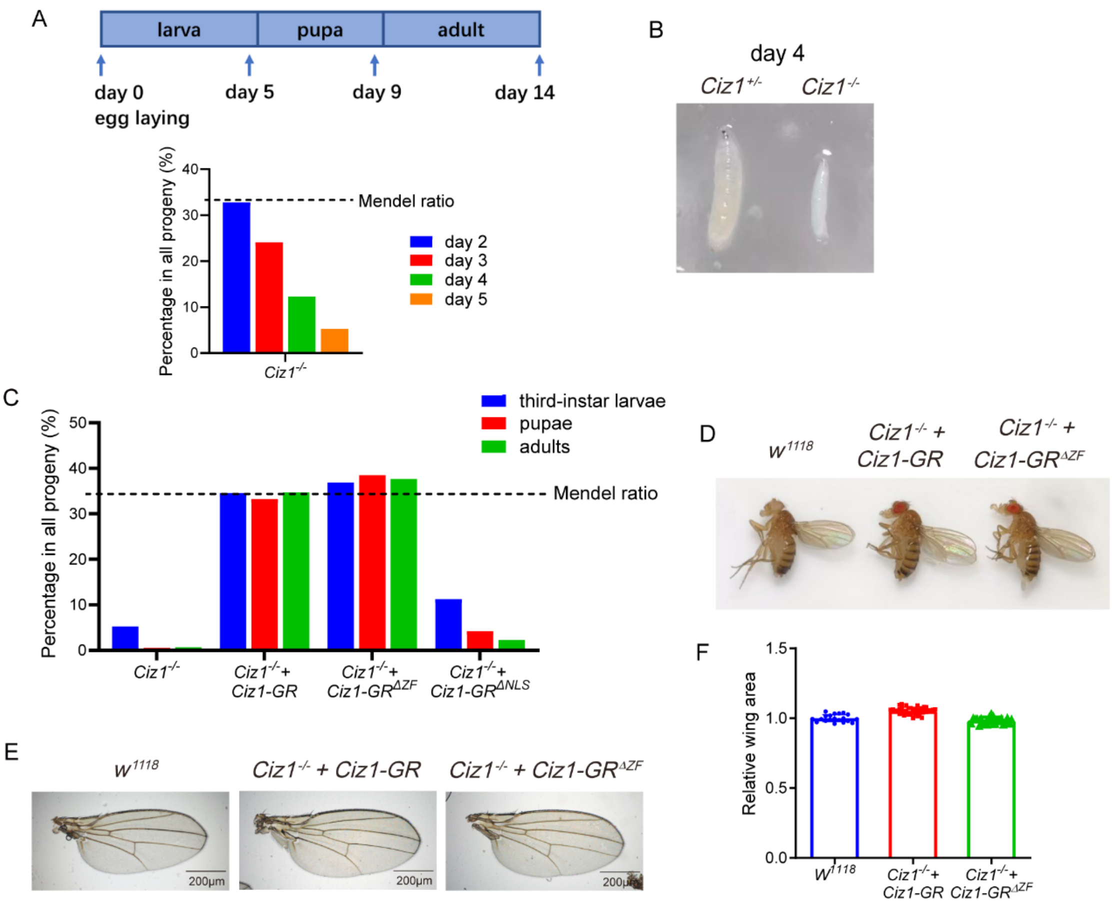


**Supplementary figure S1. *Ciz1* mutation causes larval lethality.**

(A) The percentage of *Ciz1*$^{-/-}$ animals among all progeny of the cross *Ciz1*$^{EY14316}$*/TM3,Sb,GFP* × *Ciz1*$^{KG05452}$*/ TM3,Sb,GFP* collected on the indicated time points after egg laying. The dash line shows the expected percentage of the desired genotype among all progeny of the cross. The total numbers of progeny collected on day 2, 3, 4, 5 are 464, 470, 261, 211. (B) The image of larvae on day 4 after egg laying. (C) The percentage of animals with the indicated genotypes among all progeny that survived till late third instar larvae (counted on day 5 after egg laying), pupal stage (counted on day 9 after egg laying) and adulthood (counted on day 14 after egg laying). The dash line shows the expected percentage of the desired genotype among all progeny of the cross. The total number of progeny collected from cross *Ciz1*$^{EY14316}$*/TM6B* × *Ciz1*$^{KG05452}$*/TM6B* (generating *Ciz1*$^{-/-}$), cross *Ciz1-GR/CyOGFP; Ciz1*$^{EY14316}$*/TM6B* ×

*Ciz1-GR/CyOGFP; Ciz1$^{KG05452}$/TM6B* (generating *Ciz1-GR/CyOGFP; Ciz1$^{-/-}$ or Ciz1-GR; Ciz1$^{-/-}$*), cross *Ciz1-GR$^{\Delta ZF}$/CyOGFP; Ciz1$^{EY14316}$/TM6B* × *Ciz1-GR$^{\Delta ZF}$/CyOGFP; Ciz1$^{KG05452}$/TM6B* (generating *Ciz1-GR$^{\Delta ZF}$/CyOGFP; Ciz1$^{-/-}$ or Ciz1-GR$^{\Delta ZF}$; Ciz1$^{-/-}$*) and cross *Ciz1-GR$^{\Delta NLS}$/CyOGFP; Ciz1$^{EY14316}$/TM6B* × *Ciz1-GR$^{\Delta NLS}$/CyOGFP; Ciz1$^{KG05452}$/TM6B* (generating *Ciz1-GR$^{\Delta NLS}$/CyOGFP; Ciz1$^{-/-}$ or Ciz1-GR$^{\Delta NLS}$; Ciz1$^{-/-}$*) is 289, 350, 250, and 258, respectively. (D) The representative images of adult flies showing comparable body size. The *w$^{1118}$* fly was used as control. (E) The representative images of adult wings. Scale bar, 200 μm. (F) Quantification of wing size. n= 20 (*w$^{1118}$*), 38 (*Ciz1$^{-/-}$* + *Ciz1-GR*), 24 (*Ciz1$^{-/-}$* + *Ciz1-GR$^{\Delta ZF}$*). The average wing size of *w$^{1118}$* was set as 1. The detailed genotypes of samples in this figure are listed in Supplementary table S4.

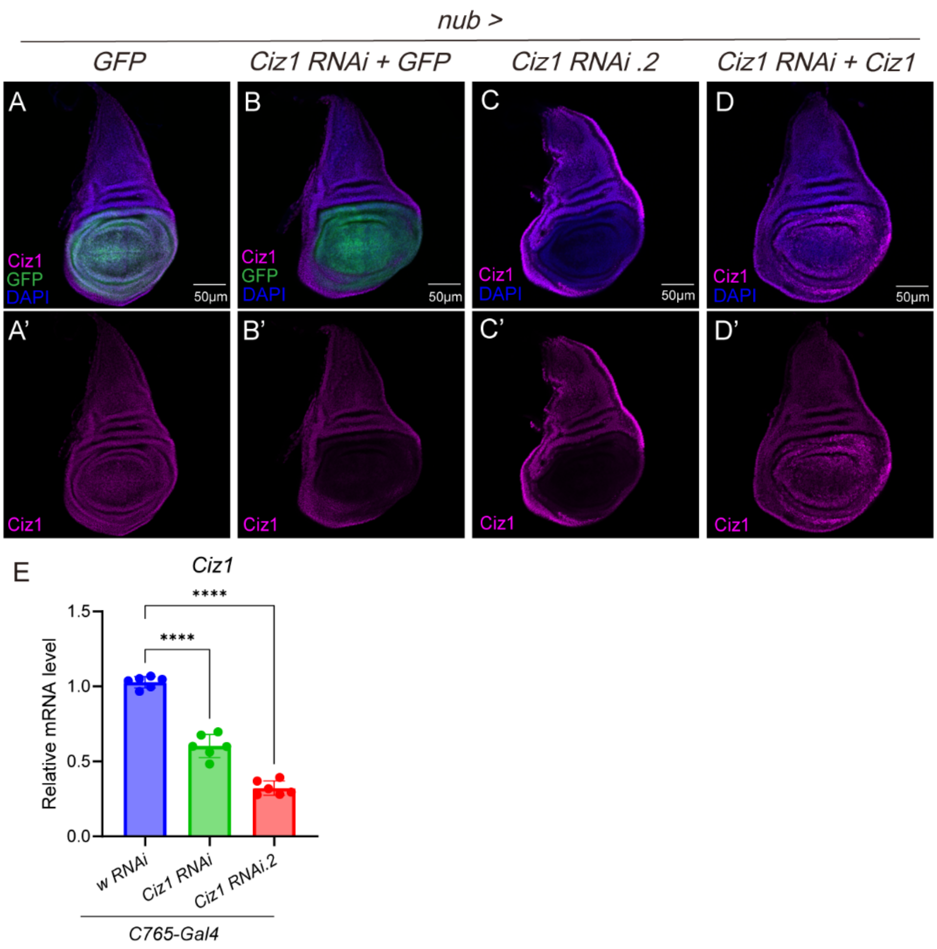


**Supplementary figure S2. The knockdown efficiency of *Ciz1 RNAi*.**

(A-D) Representative images of Ciz1 staining in wing discs of the indicated genotypes. GFP in (A) and (B) marks *nub-Gal4* expressing region. Scale bar, 50 µm. (E) RT-qPCR data showing the relative mRNA levels of Ciz1 in the indicated wing discs. n= 6. Each biological replicate was prepared from 100 discs. ****: $P< 0.0001$. The detailed genotypes of samples in this figure are listed in Supplementary table S4.

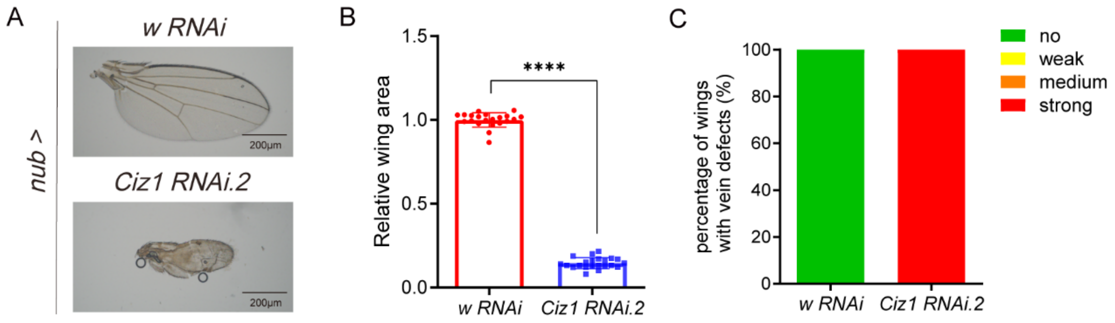


**Supplementary figure S3. Knockdown of *Ciz1* causes wing defects.**

(A) Representative images of wings with the indicated genotypes. (B) Quantification of wing sizes. The average size of *nub > w RNAi* was set as 1. n= 21 (*nub > w RNAi*) and 22 (*nub > Ciz1 RNAi.2*). ****: $P < 0.0001$. (C) The percentage of wings with vein defects. n= 21 (*nub > w RNAi*) and 32 (*nub > Ciz1 RNAi.2*). The detailed genotypes of samples in this figure are listed in Supplementary table S4.

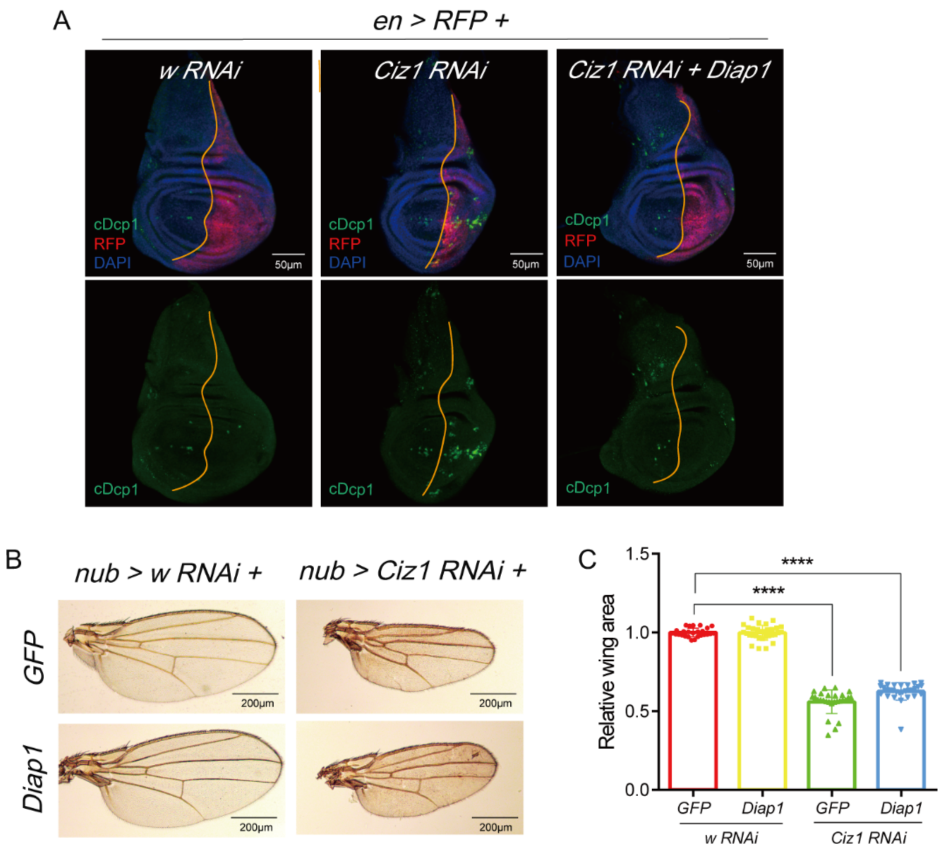


**Supplementary figure S4. The reduced wing size caused by *Ciz1* knockdown is not due to increased apoptosis.**

(A) The representative images of cDcp1 staining in wing discs with transgenes expressed in the posterior compartment labeled by RFP. The orange line marks the anterior-posterior boundary. Scale bar, 50 μm. (B) The representative images of adult wings. Scale bar, 200 μm. (C) Quantification of wing size. n= 26 (*nub> w RNAi + GFP*), 30 (*nub> w RNAi + Diap1*), 27 (*nub> Ciz1 RNAi + GFP*), 29 (*nub> Ciz1 RNAi + Diap1*). ****: *P*< 0.0001. The detailed genotypes of samples in this figure are listed in Supplementary table S4.

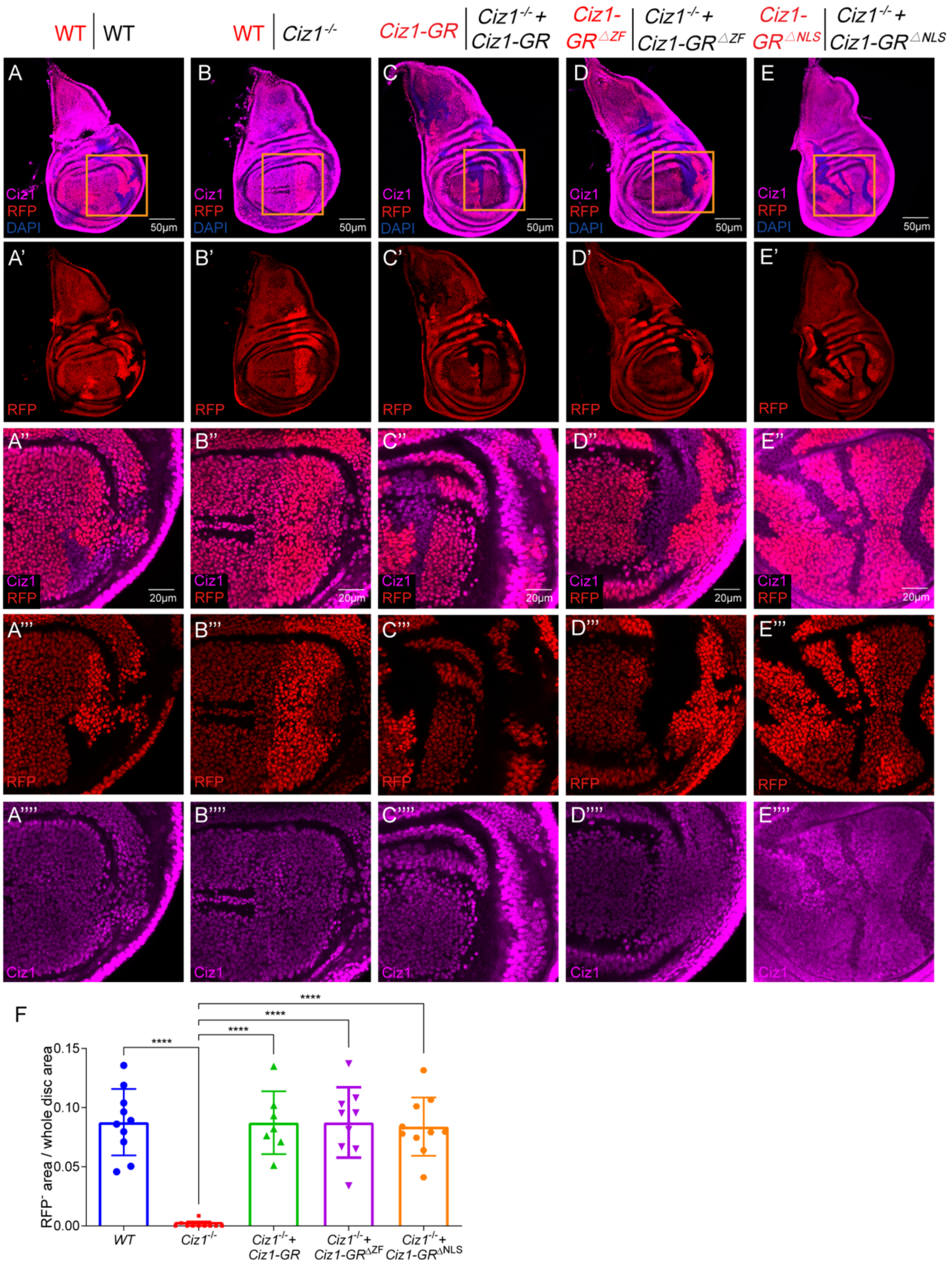


**Supplementary figure S5. *Ciz1* mutant clones in wing discs show growth disadvantage.**

(A-E) Mitotic clones in wing discs. The genotypes of RFP$^-$ and RFP$^+$ cells are shown on the top. The brighter RFP marks twin-spot clones. Images labeled with ”,’’’ and ”” were magnified images of the rectangular regions. Scale bar, 50 μm in (A-E) and (A'-E'), 20 μm in the rest images. (F) Quantification of the ratio between the $RFP^{-}$ area with the indicated genotype and the disc area. n= 10 (wild type, WT), 10 (*Ciz1*$^{-/-}$), 7 (*Ciz1*$^{-/-}$ + *Ciz1-GR*), 9 (*Ciz1*$^{-/-}$ + *Ciz1-GR*$^{\Delta ZF}$) and 10 (*Ciz1*$^{-/-}$ + *Ciz1-GR*$^{\Delta NLS}$). ****: $P< 0.0001$. The detailed genotypes of samples in this figure are listed in Supplementary table S4.

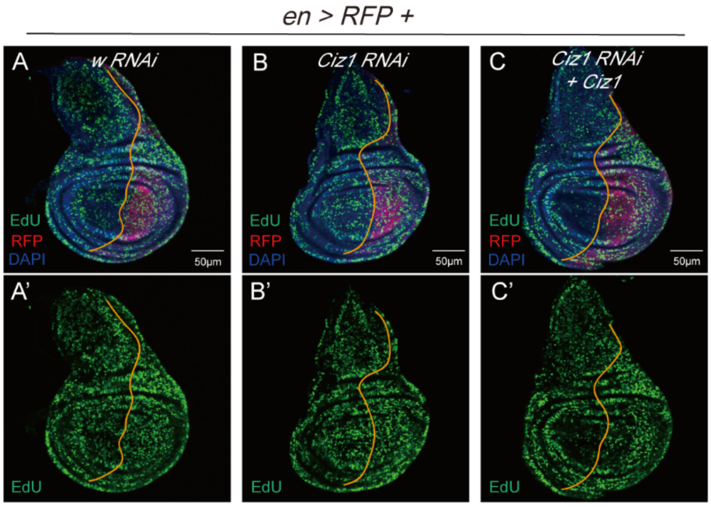


**Supplementary figure S6. Knocking down *Ciz1* does not affect DNA synthesis.**

The representative images of EdU staining in wing discs with transgenes expressed in the posterior compartment labeled by RFP. The orange line marks the anterior-posterior boundary. Scale bar, 50 µm. The detailed genotypes of samples in this figure are listed in Supplementary table S4.

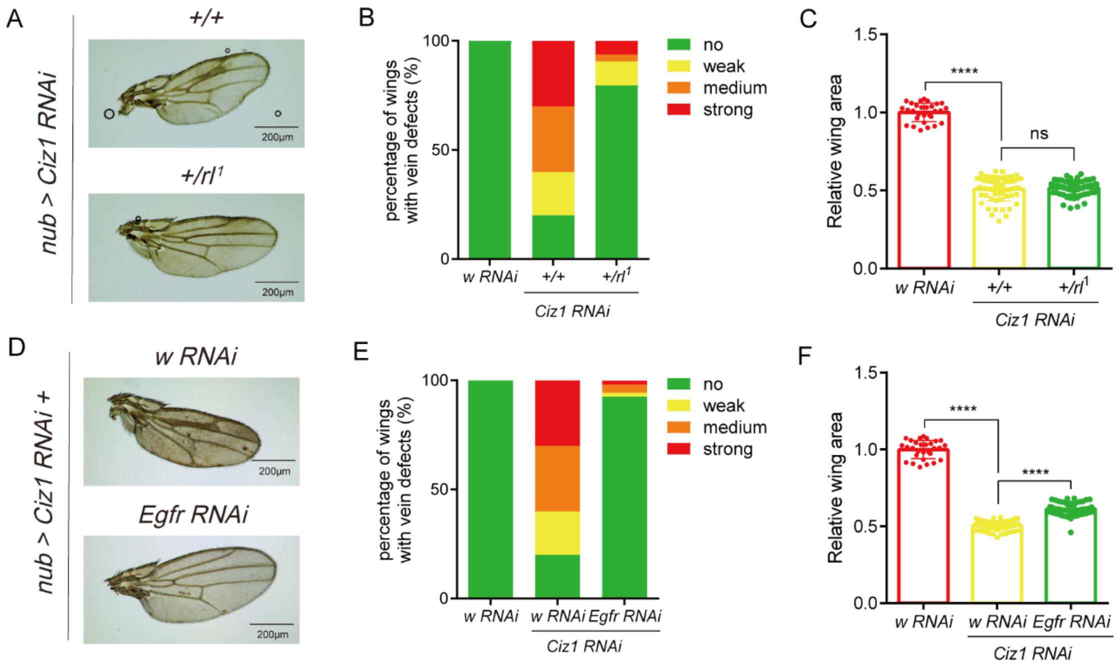


**Supplementary figure S7. Reducing EGFR-ERK signaling activity rescues vein defects caused by Ciz1 deficiency.**

(A, D) Representative images of adult wings with the indicated genotypes. Scale bar, 200 μm. (B, E) Quantification of the percentage of wings with ectopic veins. n= 31 (*nub> w RNAi*), 61 (*nub> Ciz1 RNAi*), 57 (*nub> Ciz1 RNAi,* $rl^{1/+}$) in (B) and 31 (*nub> w RNAi*), 56 (*nub> Ciz1 RNAi + w RNAi*), 54 (*nub> Ciz1 RNAi + Egfr RNAi*) in (E). (C, F) Quantification of the wing sizes. The average wing size of *nub> w RNAi* flies was set as 1. n= 29 (*nub> w RNAi*), 59 (*nub> Ciz1 RNAi*), 64 (*nub> Ciz1 RNAi,* $rl^{1/+}$) in (C) and 29 (*nub> w RNAi*), 56 (*nub> Ciz1 RNAi + w RNAi*), 50 (*nub> Ciz1 RNAi + Egfr RNAi*) in (F). ns: no significance; ****: *P*< 0.0001. The detailed genotypes of samples in this figure are listed in Supplementary table S4.

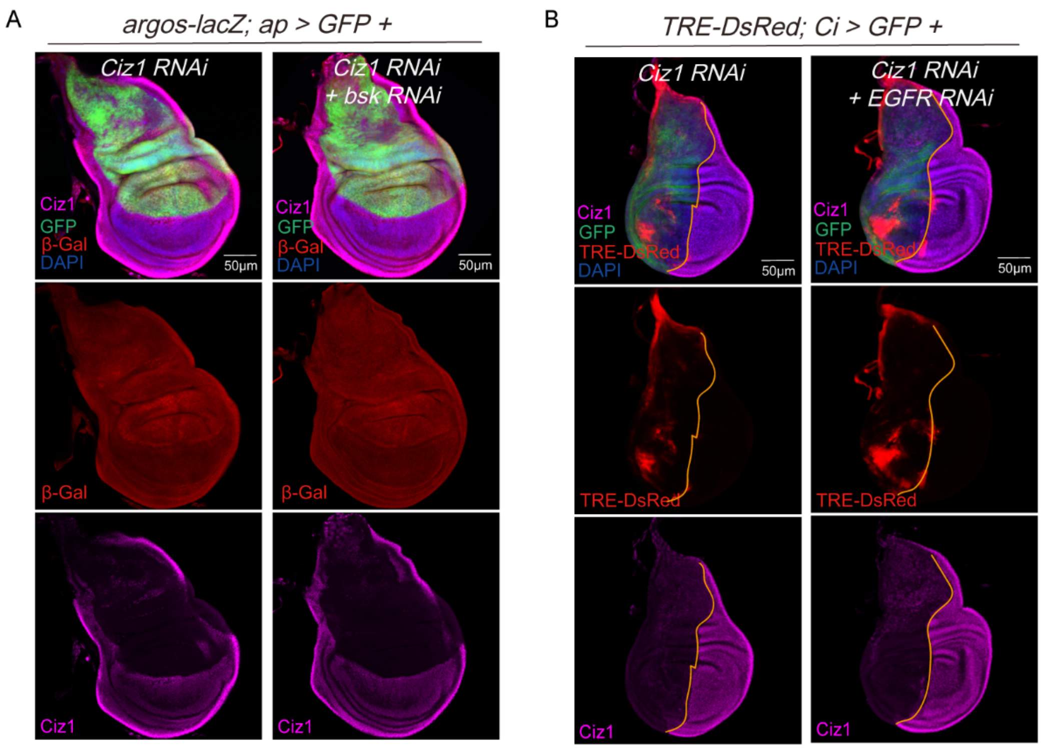


**Supplementary figure S8. Ciz1 regulates JNK and EGFR signaling in parallel.**

(A) The representative images showing *argos-lacZ* expression in wing discs with transgenes expressed in the dorsal compartment labeled by GFP. Scale bar, 50 μm. (B) The representative images showing *TRE:DsRed* expression in wing discs with transgenes expressed in the anterior compartment labeled by GFP. The orange line marks the anterior-posterior compartment. Scale bar, 50 μm. The detailed genotypes of samples in this figure are listed in Supplementary table S4.

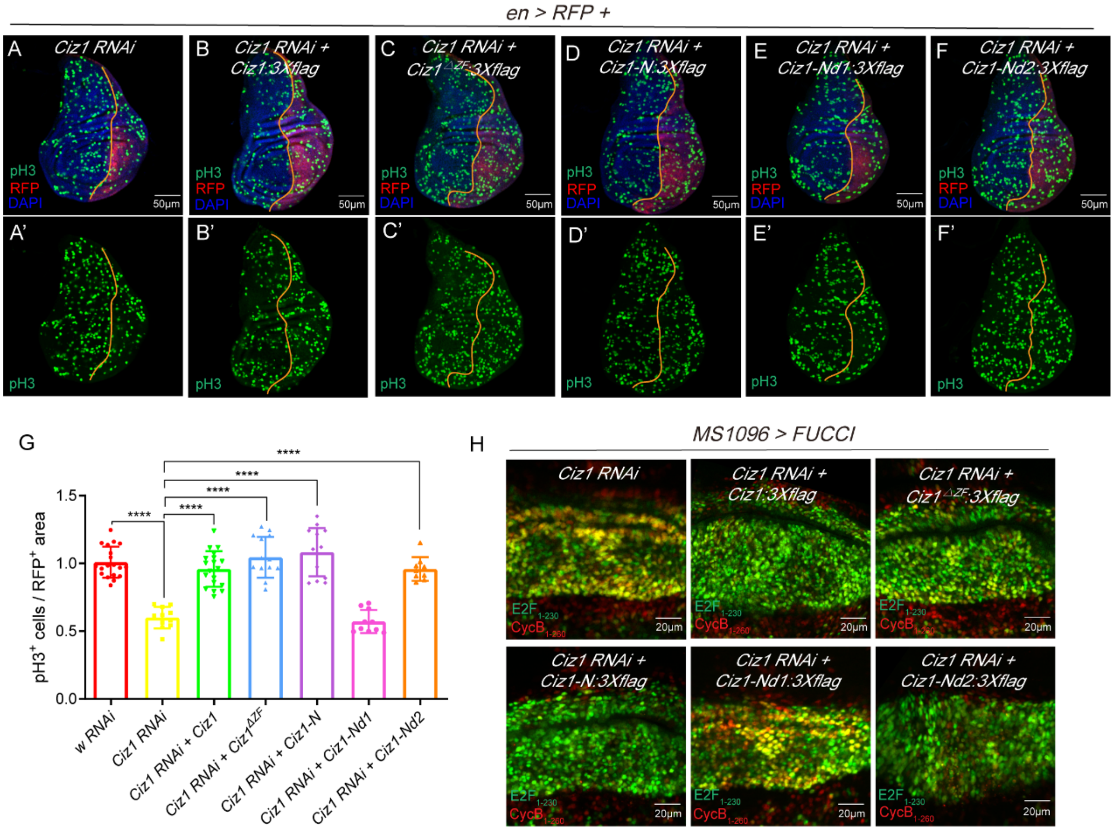


**Supplementary figure S9. The effects of different Ciz1 fragments on proliferation and cell cycle.**

(A-F) Representative images of pH3 staining in wing discs with the indicated transgenes expressed in the posterior compartment labeled by RFP. The orange line marks the anterior-posterior boundary. Scale bar, 50 µm. (G) n= 19 (*en> w RNAi*), 11 (*Ciz1 RNAi*), 19 (*Ciz1 RNAi+ Ciz1*), 14 (*Ciz1 RNAi+ Ciz1$^{\Delta ZF}$*), 13 (*Ciz1 RNAi+ Ciz1-N*), 10 (*Ciz1 RNAi+ Ciz1-Nd1*), 9 (*Ciz1 RNAi+ Ciz1-Nd2*). (H) Cell cycle analysis by FUCCI reporter in wing discs. All transgenes were expressed in the wing pouch by *MS1096-Gal4*. Scale bar, 20 µm. The detailed genotypes of samples in this figure are listed in Supplementary table S4.

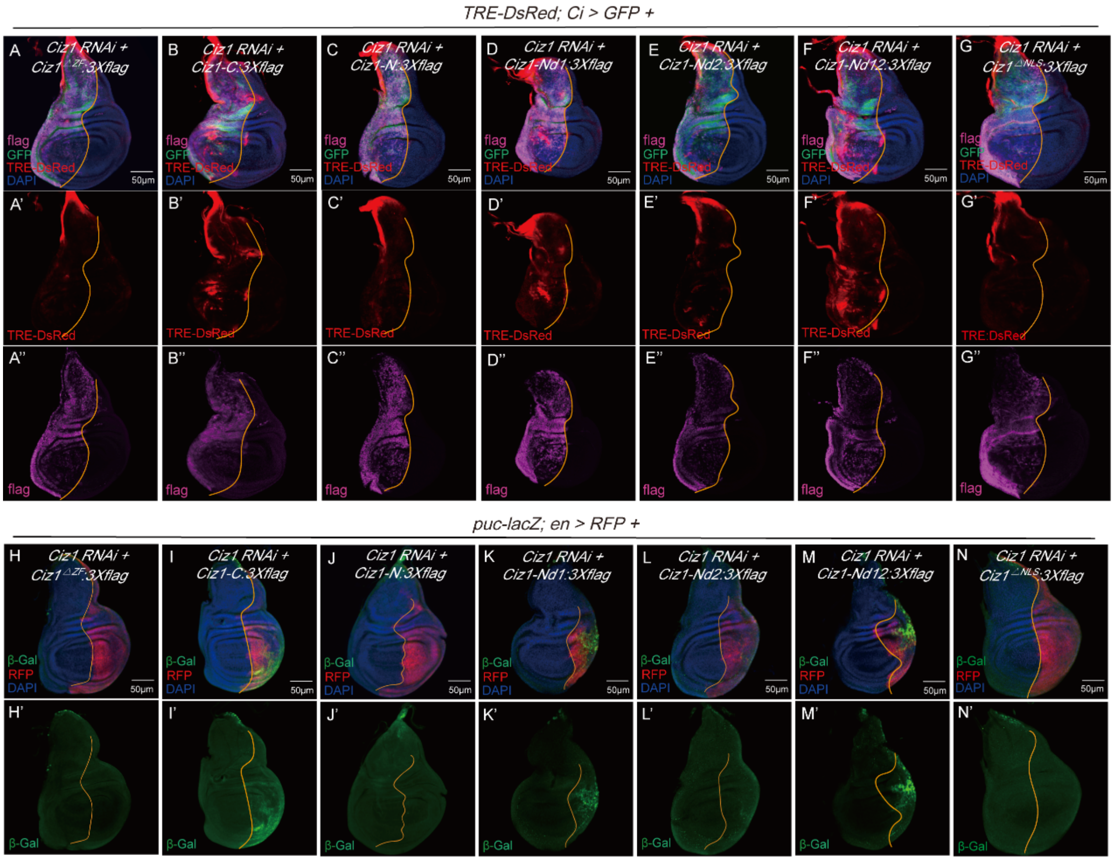


**Supplementary figure S10. The effects of different Ciz1 fragments on JNK activation.**

(A-G) The representative images showing *TRE-DsRed* expression in wing discs with the indicated transgenes expressed in the anterior compartments labeled by GFP. (H-N) The representative images showing *puc-lacZ* expression in wing discs with the indicated transgenes expressed in the posterior compartments labeled by RFP. The orange line marks the anterior-posterior boundary. Scale bar, 50 μm.

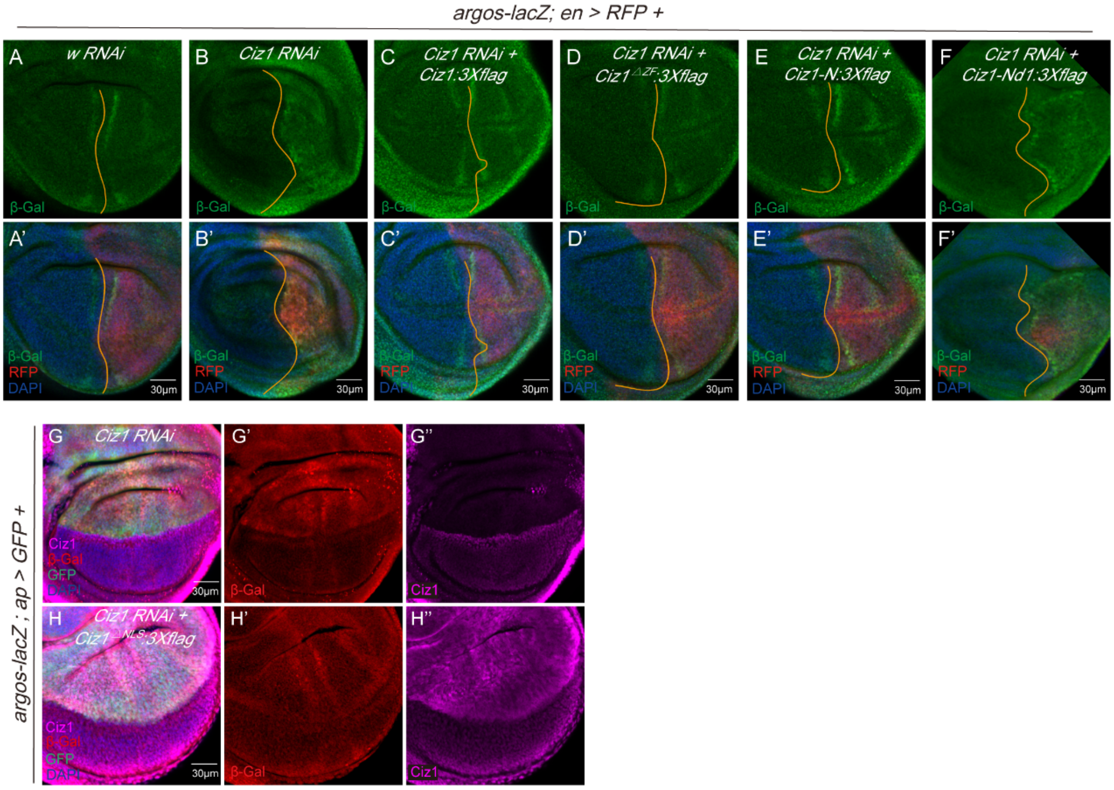


**Supplementary figure S11. The effects of different Ciz1 fragments on EGFR signaling.**

(A-F) The representative images showing *argos-lacZ* expression in wing discs with the indicated transgenes expressed in posterior compartments labeled by RFP. The orange line marks the anterior-posterior boundary. Scale bar, 30 μm. G) The representative images showing *argos-lacZ* expression in wing discs with the indicated transgenes expressed in dorsal compartments labeled by GFP. Scale bar, 30 μm.

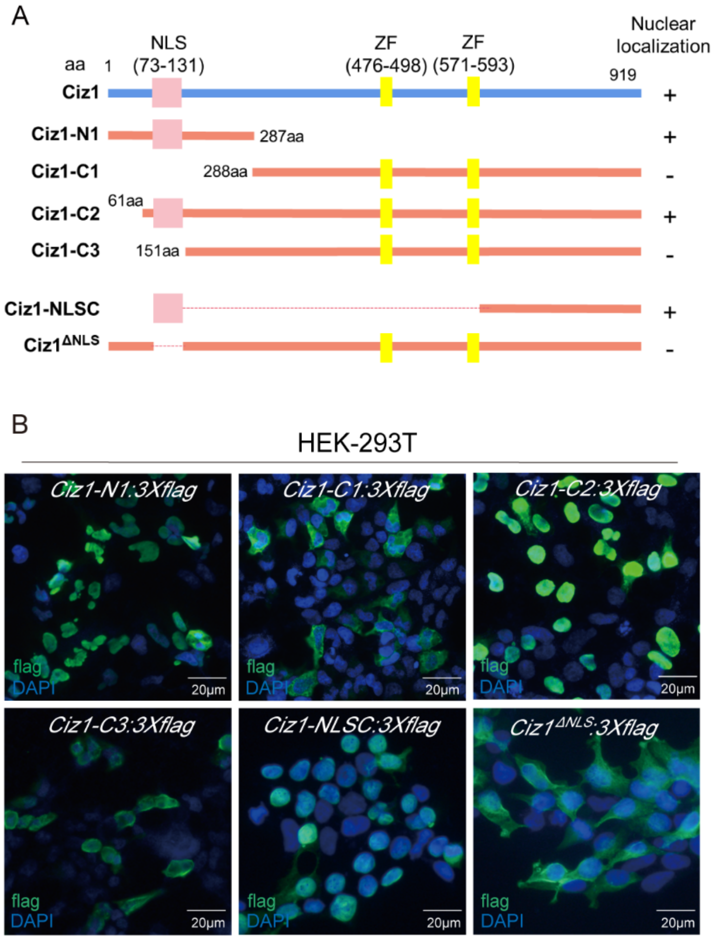


**Supplementary figure S12. Mapping the nuclear localization signal of Ciz1.**

(A) The schematic showing different Ciz1 fragments. (B) Subcellular localization of the indicated Ciz1 fragments. Scale bar, 20 µm.

**Supplementary table S1. Fly stocks used in this study.**

| Fly Stocks | Source |
|---|---|
| *UAS-w RNAi (III)* | BDSC #33623 |
| *UAS-Ciz1 RNAi.2 (II)* | VDRC #35344 |
| *UAS-Ciz1 RNAi (III)* | BDSC #27562 |
| *en-Gal4 UAS-RFP (II)* | BDSC #30557 |
| $w^{1118}$ | BCF #78 |
| *UAS-GFP:nls (II)* | BCF #80 |
| *nub-Gal4 (II)* | TB00045 |
| *tub-Gal4 (III)* | TB00129 |
| *C765-Gal4 (III)* | BDSC #36523 |
| *MS1096-Gal4 (X)* | TB00078 |
| *C311-Gal4 (II)* | BDSC #5937 |
| *UAS-Ciz1:3×flag (II)* | BDSC #92352 |
| *His2AV:mRFP FRT80B (III)* | BDSC #34499 |
| *Ciz1[EY14316] (III)* | BDSC #20919 |
| *Ciz1[KG05452] (III)* | BDSC #14595 |
| *FRT80B ry[506] (III)* | BDSC #1988 |
| *ap-Gal4 UAS-GFP (II)* | BCF #677 |
| *Ci-Gal4 UAS-GFP (III)* | BCF #444 |
| *hh-Gal4 (III)* | BCF #662 |
| *UAS-FUCCI (II)* | BDSC #55121 |
| *UAS-mito-HA-GFP(II)* | BDSC #8442 |
| *UAS-Diap1 (III)* | BDSC #6657 |
| *TRE:DsRed (II)* | BDSC #59012 |
| *puc-lacZ (III)* | BDSC #98329 |
| *argos-lacZ (III)* | BDSC #2513 |
| $rl^{1}$ | BDSC #386 |
| *UAS-rho RNAi.2 (II)* | THU5308 |
| *UAS-rho RNAi.1 (II)* | TH201501179.S |
| *UAS-bsk RNAi (II)* | TH04355.N |
| *UAS-bsk.DN (X)* | BDSC #6409 |
| *UAS-EGFR RNAi (II)* | TH201500392.S |

| | |
|---|---|
| *GstD1:GFP (II)* | BDSC #605991 |
| *UAS-Cat (II)* | BDSC #24621 |

BDSC: Bloomington Drosophila Stock Center; VDRC: Vienna Drosophila Resource Center. Stocks starting with TH and TB were obtained from Tsinghua Fly Center (Beijing, China). Stocks starting with BCF were obtained from National Drosophila Resource Center of China (NDRCC, Shanghai, China).

**Supplementary table S2. Oligonucleotides used in this study.**

| Molecular cloning primers | sequence |
|---|---|
| Ciz1/dZF/N/Nd2/N1/NLSd 1-F | gttaattaagaccggtgtggat |
| dZF 475-R-594 | ttgcgggccttgttcctaataatgagtttgatgaagctgcg |
| dZF/NLSC 594-F | attaggaacaaggcccgcaa |
| Ciz1/C/dZF/C1/C2/C2/NLSC/NLSd 919-R | tagtcaccacttccaccg |
| C 476-F | attaagaccggtgtggatcccaaaatgtgcgtccattgccgc |
| N1 287-R | cggggtacc ctcagcatcttcgccgtc |
| C1 288-F | attaagaccggtgtggatcccaaaatgaaaagcaagaagaag |
| C2 61-F | attaagaccggtgtggatcccaaaatgagcggcggaggagca |
| C3 151-F | attaagaccggtgtggatcccaaaatgaccggcggtggtggt |
| Nd1/Nd12/NLSC 73-F | ggtgtggatcccaaaatgaacaggagtcgg |
| Nd2/Nd12 131-R-165 | gtggaatcatcgcgtgggcggggtcgatcgctgcgatcat |
| NLSC 131-R-594 | tcgttgcgggccttgttcctgggtcgatcgctgcgatcat |
| NLSd 73-R-131 | ccaatgttctgcgacgaggagtcgtagtggctgttgcctc |
| NLSd 131-F | tcctcgtcgcagaacattgg |
| Nd2/Nd12 165-F | cgcccacgcgatgattccac |
| N/Nd1/Nd2/Nd12 475-R | acttccaccggtaccaatgagtttgatgaa |
| gCiz1/ gCiz1$^{\Delta ZF}$ / $^{\Delta NLS}$ -1-F | gtggccagggccgcaagcttctgtagtcgtacaaaattgaaaggc |
| gCiz1-1-R | ttacatagctgcgcttgtggct |
| gCiz1-2-1-F | agccacaagcgcagctatgtaagttggagcacttcgcactaaccta |
| gCiz1-2-1-R | agtcccgcgtttaagcttcaca |
| gCiz1-2-2-F | tgtgaagcttaaacgcgggactgagttgagcctatcgagcgttaaagc |
| gCiz1-2-2-R | cggggtcgggatatatagttggtg |
| gCiz1-3-F | caccaactatatatcccgaccccgtggcacaatgtcgatgagggg |
| gCiz1-3-R | atcctcgcttttcagagcggc |
| gCiz1-4-F | gccgctctgaaaagcgaggatgaggagactcgcaaacgcgct |
| gCiz1-4-R / gCiz1$^{\Delta ZF}$ / $^{\Delta NLS}$ -3-R | cgtcttcaagaattcgtttaaacagattgcataactgtcgaatacgaagaccc |
| gCiz1$^{\Delta ZF}$-1-R | cgcgagatgagtcctagaaacaca |
| gCiz1$^{\Delta ZF}$-2-F | tgtgtttctaggactcatctcgcg accgcagcgatgatcgcaaa |
| gCiz1$^{\Delta ZF}$-2-R | aatgagtttgatgaagctgcg |
| gCiz1$^{\Delta ZF}$-3-F | cgcagcttcatcaaactcatt attagggtgagtcagagtcattaa |

| | |
|---|---|
| gCiz1$^{\Delta NLS}$-1-R | gtcgtagtggctgttgcct |
| gCiz1$^{\Delta NLS}$-2-F | aggcaacagccactacgac tcgtcctcgtcgcagaacatt |
| gCiz1$^{\Delta NLS}$-2-R | cgctatcatccgatccggatt |
| gCiz1$^{\Delta NLS}$-3-F | aatccggatcggatgatagcg tggatgagcgcgagatagat |
| **qRT-PCR primers** | **sequence** |
| rp49-F | GCTAAGCTGTCGCACAAATG |
| rp49-R | GTTCGATCCGTAACCGATGT |
| Ciz1-F | GCCACTACGACAACAGGAGTC |
| Ciz1-R | CGCGAGATGAGTCATAGCTGC |
| rho-F | GAGCACATCTACATGCAACGC |
| rho-R | GGAGATCACTAGGATGAACCAGG |

**Supplementary table S3. Antibodies used in this study.**

| Antibodies | Company | Catalog # | Working Dilution |
|---|---|---|---|
| Ciz1 | Made in this study by Abclonal | N/A | 1:1000 |
| flag | Vazyme | RA1003 | 1:400 |
| pH3 | Cell Signaling Technology | 9706S | 1:400 |
| cDcp1 | Cell Signaling Technology | 9578S | 1:200 |
| β-Gal | SparkJade | Z3783 | 1:400 |
| anti-Rabbit Alexa Fluor 647 | ThermoFisher Scientific | A21245 | 1:200 |
| anti-Mouse Alexa Fluor 488 | ThermoFisher Scientific | A11029 | 1:200 |
| anti-Rabbit Alexa Fluor 488 | ThermoFisher Scientific | A11008 | 1:200 |
| anti-Mouse Alexa Fluor 647 | ThermoFisher Scientific | A21235 | 1:200 |
| anti-Mouse Alexa Fluor 568 | ThermoFisher Scientific | A11031 | 1:200 |

**Supplementary table S4. Genotypes of samples used in figures.**

| Panel # | Genotype |
|---|---|
| 1A | *nub-Gal4 / UAS-GFP;+/+* |
| | *nub-Gal4 / UAS-GFP; UAS-Ciz1 RNAi / +* |
| | *nub-Gal4 / UAS-Ciz1:3×flag; UAS-Ciz1 RNAi / +* |
| 2A, S6A | *en-Gal4 UAS-RFP / +; UAS-w RNAi / +* |
| 2B, S6B | *en-Gal4 UAS-RFP / +; UAS-Ciz1 RNAi / +* |
| 2C, S6C | *en-Gal4 UAS-RFP / UAS-Ciz1:3×flag; UAS-Ciz1 RNAi / +* |
| 2E | *MS1096-Gal4; UAS-FUCCI / +; UAS-w RNAi / +* |
| 2F | *MS1096-Gal4; UAS-FUCCI / +; UAS-Ciz1 RNAi / +* |
| 2G | *MS1096-Gal4; UAS-FUCCI / UAS-Ciz1:3×flag; UAS-Ciz1 RNAi / +* |
| 2H/3A-I | *+/+; UAS-w RNAi / C765-Gal4* |
| | *UAS-Ciz1 RNAi.2 / +; C765-Gal4 / +* |
| 3J | *C311-Gal4 / UAS-mito-GFP; UAS-w RNAi / +* |
| 3K | *C311-Gal4 / UAS-mito-GFP; UAS-Ciz1 RNAi / +* |
| 4A | *en-Gal4 UAS-RFP / +; UAS-w RNAi / +* |
| 4B | *en-Gal4 UAS-RFP / UAS-Ciz1 RNAi.2; + / +* |
| 4C | *GstD1:GFP / +; UAS-w RNAi / hh-Gal4* |
| 4D | *GstD1:GFP / +; UAS-Ciz1 RNAi / hh-Gal4* |
| 4E | *GstD1:GFP / UAS-Ciz1:3×flag; UAS-Ciz1 RNAi / hh-Gal4* |
| 4F | *GstD1:GFP / UAS-Cat; UAS-Ciz1 RNAi / hh-Gal4* |
| 4G | *en-Gal4 UAS-RFP / +; UAS-w RNAi / +* |
| 4H | *en-Gal4 UAS-RFP / +; UAS-Ciz1 RNAi / +* |
| 4I | *en-Gal4 UAS-RFP / UAS-Cat; UAS-Ciz1 RNAi / +* |
| 4K | *nub-Gal4 / UAS-GFP; UAS-Ciz1 RNAi / +* |
| | *nub-Gal4 / UAS-Cat; UAS-Ciz1 RNAi / +* |
| 5A | *TRE-DsRed / +; Ci-Gal4 UAS-GFP / UAS-wRNAi* |
| 5B | *TRE-DsRed / +; Ci-Gal4 UAS-GFP / UAS-Ciz1 RNAi* |
| 5C | *TRE-DsRed / UAS-Ciz1:3×flag; Ci-Gal4 UAS-GFP / UAS-Ciz1 RNAi* |
| 5D | *TRE-DsRed / UAS-Cat ; Ci-Gal4 UAS-GFP / UAS-Ciz1 RNAi* |
| 5E | *TRE-DsRed / UAS-bsk RNAi; Ci-Gal4 UAS-GFP / UAS-Ciz1 RNAi* |
| 5F | *UAS-*$bsk^{DN}$*/+; TRE-DsRed / +; Ci-Gal4 UAS-GFP / UAS-Ciz1 RNAi* |
| 5G | *en-Gal4 UAS-RFP / +; puc-lacZ / UAS-wRNAi* |

| | |
|---|---|
| 5H | *en-Gal4 UAS-RFP / +; puc-lacZ / UAS-Ciz1 RNAi* |
| 5I | *en-Gal4 UAS-RFP / UAS-Ciz1:3×flag ; puc-lacZ / UAS-Ciz1 RNAi* |
| 5J | *en-Gal4 UAS-RFP / UAS-Cat ; puc-lacZ / UAS-Ciz1 RNAi* |
| 5K | *en-Gal4 UAS-RFP / UAS-bsk RNAi; puc-lacZ / UAS-Ciz1 RNAi* |
| 5L | *UAS-bsk$^{DN}$; en-Gal4 UAS-RFP / +; puc-lacZ / UAS-Ciz1 RNAi* |
| 5M | *en-Gal4 UAS-RFP / +; UAS-w RNAi / +* |
| 5N | *en-Gal4 UAS-RFP / +; UAS-Ciz1 RNAi / +* |
| 5O | *en-Gal4 UAS-RFP / UAS-bsk RNAi; UAS-Ciz1 RNAi / +* |
| 5P | *UAS-bsk$^{DN}$; en-Gal4 UAS-RFP / +; UAS-Ciz1 RNAi / +* |
| 6A | *ap-Gal4 UAS-GFP / +; UAS-w RNAi / argos-lacZ* |
| 6B | *ap-Gal4 UAS-GFP / +; UAS-Ciz1 RNAi / argos-lacZ* |
| 6C | *ap-Gal4 UAS-GFP / UAS-Ciz1:3×flag; UAS-Ciz1 RNAi / argos-lacZ* |
| 6D | *ap-Gal4 UAS-GFP / UAS-Cat; UAS-Ciz1 RNAi / argos-lacZ* |
| 6E | *ap-Gal4 UAS-GFP / UAS-rho RNAi.1; UAS-Ciz1 RNAi / argos-lacZ* |
| 6F | *tub-Gal4 / +; UAS-w RNAi / +* |
| | *tub-Gal4 / +; UAS-Ciz1 RNAi / +* |
| 6G | *nub-Gal4 / +; UAS-w RNAi / +* |
| | *nub-Gal4 / +; UAS-Ciz1 RNAi / UAS-w RNAi* |
| | *nub-Gal4 / UAS-rho RNAi.1; UAS-Ciz1 RNAi / +* |
| | *nub-Gal4 / UAS-rho RNAi.2; UAS-Ciz1 RNAi / +* |
| 7C | *nub-Gal4 / UAS-GFP; UAS-Ciz1 RNAi / +* |
| | *nub-Gal4 / UAS-Ciz1:3×flag; UAS-Ciz1 RNAi / +* |
| | *nub-Gal4 / UAS-Ciz1$^{\Delta ZF}$:3×flag; UAS-Ciz1 RNAi / +* |
| | *nub-Gal4 / UAS-Ciz1-C:3×flag; UAS-Ciz1 RNAi / +* |
| | *nub-Gal4 / UAS-Ciz1-N:3×flag; UAS-Ciz1 RNAi / +* |
| | *nub-Gal4 / UAS-Ciz1-Nd1:3×flag; UAS-Ciz1 RNAi / +* |
| | *nub-Gal4 / UAS-Ciz1-Nd2:3×flag; UAS-Ciz1 RNAi / +* |
| | *nub-Gal4 / UAS-Ciz1-Nd12:3×flag; UAS-Ciz1 RNAi / +* |
| 7E | *GstD1:GFP / +; UAS-Ciz1 RNAi / hh-Gal4* |
| 7F | *GstD1:GFP / UAS-Ciz1$^{\Delta ZF}$:3×flag; UAS-Ciz1 RNAi / hh-Gal4* |
| 7G | *GstD1:GFP / UAS-Ciz1-N:3×flag; UAS-Ciz1 RNAi / hh-Gal4* |
| 7H | *GstD1:GFP / UAS-Ciz1-Nd1:3×flag; UAS-Ciz1 RNAi / hh-Gal4* |
| 7I | *GstD1:GFP / UAS-Ciz1-Nd2:3×flag; UAS-Ciz1 RNAi / hh-Gal4* |
| 7J | *GstD1:GFP / UAS-Ciz1-Nd12:3×flag; UAS-Ciz1 RNAi / hh-Gal4* |

| | |
|---|---|
| 8A | *w*$^{1118}$ |
| 8B | *en-Gal4 UAS-RFP / UAS-Ciz1:3×flag* |
| 8C | *en-Gal4 UAS-RFP / UAS-Ciz1-N:3×flag* |
| 8D | *en-Gal4 UAS-RFP / UAS-Ciz1-C:3×flag* |
| 8E | *en-Gal4 UAS-RFP / UAS-Ciz1*$^{\Delta NLS}$*:3×flag* |
| 8F | *en-Gal4 UAS-RFP / UAS-Ciz1-NLSC:3×flag* |
| 8G | *GstD1:GFP / +; UAS-Ciz1 RNAi / hh-Gal4* |
| 8H | *GstD1:GFP / UAS-Ciz1*$^{\Delta NLS}$*:3×flag; UAS-Ciz1 RNAi / hh-Gal4* |
| 8I | *en-Gal4 UAS-RFP / +; UAS-Ciz1 RNAi / +* |
| 8J | *en-Gal4 UAS-RFP / UAS-Ciz1*$^{\Delta NLS}$*:3×flag; UAS-Ciz1 RNAi / +* |
| 8L | *nub-Gal4 / +; UAS-Ciz1 RNAi / UAS- w RNAi* |
| | *nub-Gal4 / UAS-Ciz1*$^{\Delta NLS}$*:3×flag; UAS-Ciz1 RNAi / +* |
| S1A | *Ciz1*$^{EY14316}$ */ Ciz1*$^{KG05452}$ |
| S1B | *Ciz1*$^{EY14316}$*/ TM3,Sb,GFP or Ciz1* $^{KG05452}$ */ TM3,Sb,GFP* |
| | *Ciz1*$^{EY14316}$ */ Ciz1*$^{KG05452}$ |
| S1C-F | *w*$^{1118}$ |
| | *Ciz1*$^{EY14316}$ */ Ciz1*$^{KG05452}$ |
| | *Ciz1-GR / Ciz1-GR; Ciz1*$^{EY14316}$ */ Ciz1*$^{KG05452}$ or *Ciz1-GR / +; Ciz1*$^{EY14316}$ */ Ciz1*$^{KG05452}$ |
| | *Ciz1-GR*$^{\Delta ZF}$ */ Ciz1-GR*$^{\Delta ZF}$*; Ciz1*$^{EY14316}$ */ Ciz1*$^{KG05452}$ or *Ciz1-GR*$^{\Delta ZF}$ */ +; Ciz1*$^{EY14316}$ */ Ciz1*$^{KG05452}$ |
| | *Ciz1-GR*$^{\Delta NLS}$ */ Ciz1-GR*$^{\Delta NLS}$*; Ciz1*$^{EY14316}$ */ Ciz1*$^{KG05452}$ or *Ciz1-GR*$^{\Delta NLS}$ */ +; Ciz1*$^{EY14316}$ */ Ciz1*$^{KG05452}$ |
| S2A | *nub-Gal4 / UAS-GFP; UAS- w RNAi / +* |
| S2B | *nub-Gal4 / UAS-GFP; UAS-Ciz1 RNAi / +* |
| S2C | *nub-Gal4 / UAS-Ciz1 RNAi.2; + / +* |
| S2D | *nub-Gal4 / UAS-Ciz1:3×flag; UAS-Ciz1 RNAi / +* |
| S2E | *+/+; UAS-w RNAi / C765-Gal4* |
| | *+/+; UAS-Ciz1 RNAi / C765-Gal4* |
| | *UAS-Ciz1 RNAi.2 / +; C765-Gal4 / +* |
| S3A | *nub-Gal4 / +; UAS-w RNAi / +* |
| | *nub-Gal4 / UAS-Ciz1 RNAi.2; + / +* |
| S4A | *en-Gal4 UAS-RFP / +; UAS-w RNAi / +* |
| | *en-Gal4 UAS-RFP / +; UAS-Ciz1 RNAi / +* |

| | |
|---|---|
| | *en-Gal4 UAS-RFP / +; UAS-Ciz1 RNAi / UAS-Diap1* |
| S4B | *nub-Gal4 / UAS-GFP; UAS-w RNAi / +* |
| | *nub-Gal4 / UAS-GFP; UAS-Ciz1 RNAi / +* |
| | *nub-Gal4 / +; UAS-Diap1 / UAS-w RNAi* |
| | *nub-Gal4 / +; UAS-Diap1 / UAS-Ciz1 RNAi* |
| S5A | *hsflp; His2AVmRFP FRT80B / FRT80B* |
| S5B | *hsflp; His2AVmRFP FRT80B / Ciz1*$^{EY14316}$ *FRT80B* |
| S5C | *hsflp; Ciz1-GR / +; His2AVmRFP FRT80B / Ciz1*$^{EY14316}$ *FRT80B* |
| S5D | *hsflp; Ciz1-GR*$^{\Delta ZF}$ */ +; His2AVmRFP FRT80B / Ciz1*$^{EY14316}$ *FRT80B* |
| S5E | *hsflp; Ciz1-GR*$^{\Delta NLS}$ */ +; His2AVmRFP FRT80B / Ciz1*$^{EY14316}$ *FRT80B* |
| S7A | *w*$^{1118}$*; nub-Gal4 / +; UAS-Ciz1 RNAi / +* |
| | *rl*$^{1}$*/nub-Gal4; UAS-Ciz1 RNAi / +* |
| S7D | *nub-Gal4 / +; UAS-Ciz1 RNAi / UAS- w RNAi* |
| | *nub-Gal4 / UAS- EGFR RNAi; UAS-Ciz1 RNAi* |
| S8A | *ap-Gal4 UAS-GFP / +; UAS-Ciz1 RNAi / argos-lacZ* |
| | *ap-Gal4 UAS-GFP / UAS-bsk RNAi; UAS-Ciz1 RNAi / argos-lacZ* |
| S8B | *TRE-DsRed / +; Ci-Gal4 UAS-GFP / UAS-Ciz1 RNAi* |
| | *TRE-DsRed / UAS-EGFR RNAi; Ci-Gal4 UAS-GFP / UAS-Ciz1 RNAi* |
| S9A | *en-Gal4 UAS-RFP / +; UAS-Ciz1 RNAi / +* |
| S9B | *en-Gal4 UAS-RFP / UAS-Ciz1:3×flag; UAS-Ciz1 RNAi / +* |
| S9C | *en-Gal4 UAS-RFP / UAS-Ciz1*$^{\Delta ZF}$*:3×flag; UAS-Ciz1 RNAi / +* |
| S9D | *en-Gal4 UAS-RFP / UAS-Ciz1-N:3×flag; UAS-Ciz1 RNAi / +* |
| S9E | *en-Gal4 UAS-RFP / UAS-Ciz1-Nd1:3×flag; UAS-Ciz1 RNAi / +* |
| S9F | *en-Gal4 UAS-RFP / UAS-Ciz1-Nd2:3×flag; UAS-Ciz1 RNAi / +* |
| S9H | *MS1096-Gal4/+; UAS-FUCCI / +; UAS-Ciz1 RNAi / +* |
| | *MS1096-Gal4/+; UAS-FUCCI / UAS-Ciz1:3×flag; UAS-Ciz1 RNAi /+* |
| | *MS1096-Gal4/+; UAS-FUCCI / UAS-Ciz1*$^{\Delta ZF}$*:3×flag; UAS-Ciz1 RNAi /+* |
| | *MS1096-Gal4/+; UAS-FUCCI / UAS-Ciz1-N:3×flag; UAS-Ciz1 RNAi / +* |
| | *MS1096-Gal4/+; UAS-FUCCI/UAS-Ciz1-Nd1:3×flag; UAS-Ciz1 RNAi/+* |
| | *MS1096-Gal4/+; UAS-FUCCI/UAS-Ciz1-Nd2:3×flag; UAS-Ciz1 RNAi/+* |
| S10A | *TRE-DsRed / UAS-Ciz1*$^{\Delta ZF}$*:3×flag; Ci-Gal4 UAS-GFP / UAS-Ciz1 RNAi* |

| | |
|---|---|
| S10B | *TRE-DsRed / UAS-Ciz1-C:3×flag; Ci-Gal4 UAS-GFP / UAS-Ciz1 RNAi* |
| S10C | *TRE-DsRed / UAS-Ciz1-N:3×flag; Ci-Gal4 UAS-GFP / UAS-Ciz1 RNAi* |
| S10D | *TRE-DsRed / UAS-Ciz1-Nd1:3×flag; Ci-Gal4 UAS-GFP / UAS-Ciz1 RNAi* |
| S10E | *TRE-DsRed / UAS-Ciz1-Nd2:3×flag; Ci-Gal4 UAS-GFP / UAS-Ciz1 RNAi* |
| S10F | *TRE-DsRed / UAS-Ciz1-Nd12:3×flag; Ci-Gal4 UAS-GFP / UAS-Ciz1 RNAi* |
| S10G | *TRE-DsRed / UAS-*$Ciz1^{\Delta NLS}$*:3×flag; Ci-Gal4 UAS-GFP / UAS-Ciz1 RNAi* |
| S10H | *en-Gal4 UAS-RFP / UAS-*$Ciz1^{\Delta ZF}$*:3×flag; puc-lacZ / UAS-Ciz1 RNAi* |
| S10I | *en-Gal4 UAS-RFP / UAS-Ciz1-C:3×flag; puc-lacZ / UAS-Ciz1 RNAi* |
| S10J | *en-Gal4 UAS-RFP / UAS-Ciz1-N:3×flag; puc-lacZ / UAS-Ciz1 RNAi* |
| S10K | *en-Gal4 UAS-RFP / UAS-Ciz1-Nd1:3×flag; puc-lacZ / UAS-Ciz1 RNAi* |
| S10L | *en-Gal4 UAS-RFP / UAS-Ciz1-Nd2:3×flag; puc-lacZ / UAS-Ciz1 RNAi* |
| S10M | *en-Gal4 UAS-RFP / UAS-Ciz1-Nd12:3×flag; puc-lacZ / UAS-Ciz1 RNAi* |
| S10N | *en-Gal4 UAS-RFP / UAS-*$Ciz1^{\Delta NLS}$*:3×flag; puc-lacZ / UAS-Ciz1 RNAi* |
| S11A | *en-Gal4 UAS-RFP / +; UAS-w RNAi / argos-lacZ* |
| S11B | *en-Gal4 UAS-RFP / +; UAS-Ciz1 RNAi / argos-lacZ* |
| S11C | *en-Gal4 UAS-RFP / UAS-Ciz1:3×flag; UAS-Ciz1 RNAi / argos-lacZ* |
| S11D | *en-Gal4 UAS-RFP / UAS-*$Ciz1^{\Delta ZF}$*:3×flag; UAS-Ciz1 RNAi / argos-lacZ* |
| S11E | *en-Gal4 UAS-RFP / UAS-Ciz1-N:3×flag; UAS-Ciz1 RNAi / argos-lacZ* |
| S11F | *en-Gal4 UAS-RFP / UAS-Ciz1-Nd1:3×flag; UAS-Ciz1 RNAi / argos-lacZ* |
| S11G | *ap-Gal4 UAS-GFP / +; UAS-Ciz1 RNAi / argos-lacZ* |
| S11H | *ap-Gal4 UAS-GFP / UAS-*$Ciz1^{\Delta NLS}$*:3×flag; UAS-Ciz1 RNAi / argos-lacZ* |
| S12B | *pCDNA3-Ciz1-N1:3×flag* |
| | *pCDNA3-Ciz1-C1:3×flag* |
| | *pCDNA3-Ciz1-C2:3×flag* |
| | *pCDNA3-Ciz1-C3:3×flag* |
| | *pCDNA3-Ciz1-NLSC:3×flag* |
| | *pCDNA3-*$Ciz1^{\Delta NLS}$*:3×flag* |
